# A gene regulatory element modulates myosin expression and controls cardiomyocyte response to stress

**DOI:** 10.1101/2025.07.19.665672

**Authors:** Taylor Anglen, Irene M. Kaplow, Baekgyu Choi, Enya Dewars, Robin M. Perelli, Kevin T. Hagy, Duc Tran, Megan E. Ramaker, Svati Shah, Inkyung Jung, Andrew P. Landstrom, Ravi Karra, Yarui Diao, Charles A. Gersbach

## Abstract

A hallmark of heart disease is gene dysregulation and reactivation of fetal gene programs. Reactivation of these fetal programs has compensatory effects during heart failure, depending on the type and stage of the underlying cardiomyopathy. Thousands of putative cardiac gene regulatory elements have been identified that may control these programs, but their functions are largely unknown. We profile genome-wide changes to gene expression and chromatin structure in cardiomyocytes derived from human pluripotent stem cells. We identify and characterize a gene regulatory element essential for the regulation of *MYH6*, which encodes human fetal myosin. Using chromatin conformation assays in combination with epigenome editing, we find that gene regulation is mediated by direct interaction between *MYH6* and the enhancer. We also find that enhancer activation alters cardiomyocyte response to the hypertrophy-inducing peptide endothelin-1. Enhancer activation prevents polyploidization and changes in calcium dynamics following stress with endothelin-1. Collectively, these results identify regulatory mechanisms of cardiac gene expression programs that modulate cardiomyocyte maturation, cellular stress response, and could serve as potential therapeutic targets.

## Introduction

Cardiac development is a tightly controlled process driven by temporal and spatial coordination of gene expression. Critical transcription factors, signaling pathways, and genes important for proper cardiac development have been identified. Disruption of these key components has been implicated in various diseases at all stages of life, ranging from congenital heart disease to heart failure (Yuan et al. 2021; Dirkx et al. 2013; May et al. 2012).

Cardiac gene regulation is typically coordinated by multiple distal *cis*-regulatory elements (CREs), many of which also change activity with stress (Man et al. 2021; Sergeeva et al. 2016a) Thousands of putative CREs have been identified in cardiac tissue. Reporter assays have shown that CREs can be chamber-specific or vary in activity across each chamber of the heart. Many of these putative CREs function as enhancers – CREs distal to transcription start sites that regulate the cell-type-specificity of gene expression. However, few enhancers have been evaluated for their endogenous function, their target gene, or their roles in conditions of stress (May et al. 2012).

Interestingly, many of these key developmental factors are dysregulated in cardiomyocytes (CMs) during heart failure and following heart injury to compensate for the increased demand on the heart. The change in CM transcription is driven, in large part, by a response to paracrine signals (Talman and Kivelä 2018; Man et al. 2021; Jiang et al. 2019). Specific alterations are temporarily beneficial but eventually become maladaptive (Wende et al. 2017; Kerkelä et al. 2015a). Others are cardioprotective, correlate with better patient outcomes, and are potential therapeutic targets (Goetze et al. 2020; Kerkelä et al. 2015a). Genome-wide epigenetic modulation using HDAC inhibitors can be beneficial during heart failure, implicating gene regulation as a therapeutic target (Gallo et al. 2008).

To better understand the function of the noncoding regulatory genome in CMs, we performed ATAC-sequencing and RNA-sequencing on human induced pluripotent stem cell-derived cardiomyocytes (iPSC-CMs). From these data, we identified CREs in proximity to differentially expressed genes, one of which is proximal to the *MYH6* and *MYH7* genes that encode cardiac myosin isoforms. We found this enhancer to be important for regulating myosin expression and the switch between the human fetal isoform, *MYH6*, and mature isoform, *MYH7*. *MYH6* is associated with a spectrum of disease-related phenotypes, highlighting the importance of studying its regulation. *MYH6* expression is maintained at low levels into adulthood but is silenced during heart failure, leading to slower contractions of the heart (Chen et al. 2021; Lowes et al. 1997; Nakao et al. 1997; Miyata et al. 2000). Activation has been proposed as a potential treatment for DCM and end-stage heart failure (Locher et al. 2011; Nakao et al. 1997). Overexpression of MYH6 improved contractile force generation in cultured primary CMs from failing human and rabbit hearts (Locher et al. 2011; Herron et al. 2010). Mutations in *MYH6* cause congenital heart defects and severe forms of dilated cardiomyopathy (DCM) and hypertrophic cardiomyopathy (HCM) (Carniel et al. 2005; Posch et al. 2011). Knock-down of MYH6 in chick embryos resulted in atrial septal defects during heart development (Ching et al. 2005)Despite years of work investigating the consequences of *MYH6* silencing and loss-of-function mutations, the regulatory mechanism of *MYH6* expression is not fully understood. An increased understanding of this mechanism could inform the development of future therapeutics.

We report an enhancer near *MYH6* and *MYH7* that is functional in human iPSC-CMs, as well as its mouse ortholog that is functional in a mouse cardiac cell line, and show that this enhancer modulates the cellular response to the pro-hypertrophic GPCR agonist Endothelin-1 (ET-1) as a model of cell stress. Using CRISPR epigenome editing and targeted chromatin conformation analysis, we demonstrate that direct contact between this enhancer and *MYH6* is necessary for *MYH6* expression and that toggling the activity of the enhancer results in the alteration of chromatin looping and myosin gene expression. The role of this enhancer in developmental states is further supported by epigenetic signatures across chambers of the heart (Wei et al. 2022; Yang et al. 2018; Locher et al. 2011). We observed that cell stress leads to the loss of proximal interactions surrounding *MYH6* in iPSC-CMs and that enhancer activation significantly reduces this loss of physical contact. Finally, we see that enhancer activation prevents reduced amplitude of calcium signaling and reduces the increase in genomic content that occurs after culture with ET-1. These findings support a new mechanism of myosin gene regulation and implicate myosin regulation as directly influencing cardiomyocyte response to stress.

## Results

### WNT activation promotes immature phenotypes in cultured iPSC-CMs

To identify enhancers that regulate immature gene expression, we aimed to develop a reproducible and inducible method to drive immature phenotypes in human iPSC-CMs. We chose to activate WNT signaling through chemical inhibition of GSK3 (GSK3i) to induce the immature state in iPSC-CMs (Singh et al. 2019; Mills et al. 2019; Buikema et al. 2020; Mollova et al. 2013). GSK3 inhibition promotes immature phenotypes, including increased markers of cell cycling such as DNA synthesis and mitosis, reduced cell ploidy, and sarcomere disassembly (Singh et al. 2019; Mills et al. 2019; Buikema et al. 2020; Mollova et al. 2013).

CHIR99021 (CHIR) is a potent GSK3 inhibitor that promotes CM proliferation (Singh et al. 2019; Buikema et al. 2020; Mills et al. 2019). We treated cells with 0 µM, 2 µM, or 4 µM CHIR following iPSC to CM differentiation and 1 week of culture (**Fig. 1a, Supplemental Table S1**). We observed sarcomere disassembly following GSK3 inhibition, consistent with previous reports (**Fig. 1b**) (Buikema et al. 2020; Mills et al. 2019). GSK3 inhibition also resulted in a dose-dependent increase in markers of cardiac immaturity, including a higher frequency of cell cycle markers for S-phase entry and DNA synthesis, indicated by EdU (5-ethynyl-2’-deoxyuridine) incorporation, as well as the mitotic marker pHH3 (Cavanagh et al. 2011; Singh et al. 2019; Mills et al. 2019; Buikema et al. 2020). At baseline, 14.9% of iPSC-CMs were EdU-positive after 48 hours of culture with EdU. The addition of 2 µM and 4 µM CHIR to the media increased this to 32.0% and 40.1% EdU-positive cells, respectively (**Fig. 1b-c**). GSK3 inhibition also increased the fraction of phospho-histone H3 (pHH3)-positive nuclei from 0.03% to 0.33% and 0.38% with 2 µM and 4 µM CHIR, respectively (**Fig. 1d**) (Mills et al. 2019; Mollova et al. 2013).

**Figure 1.**
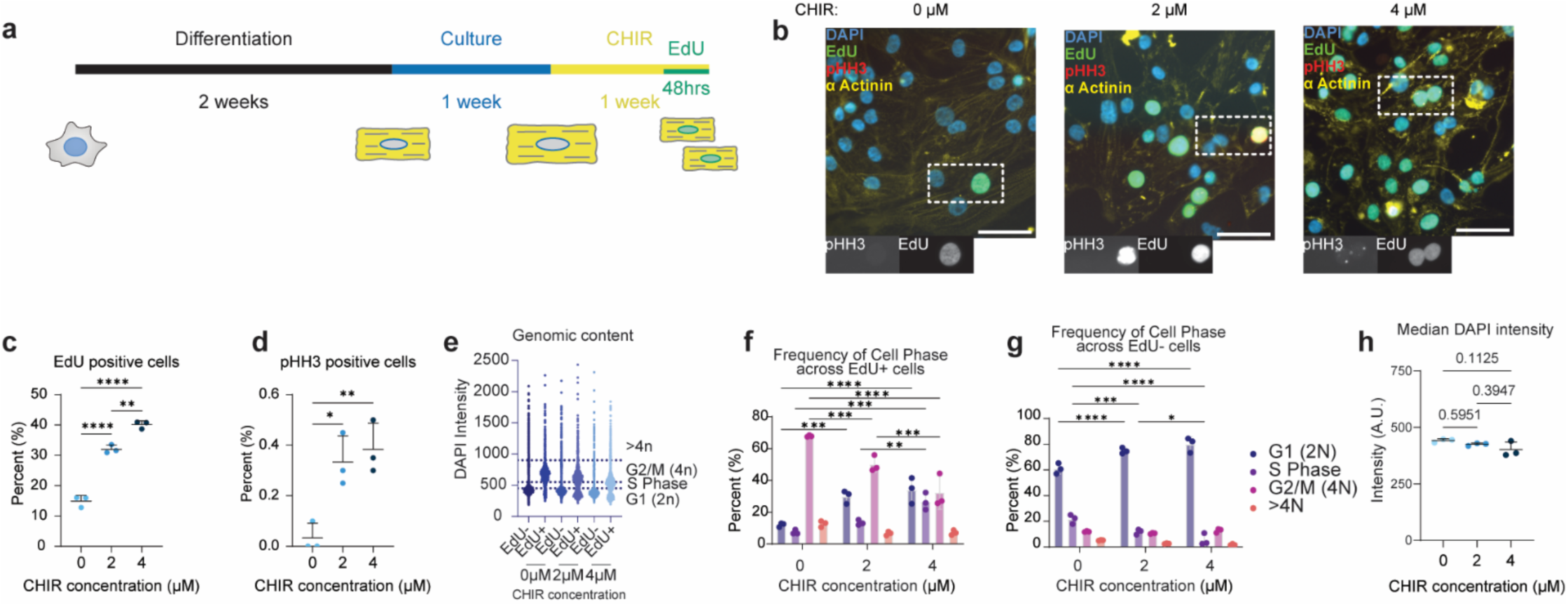
WNT activation promotes immature phenotypes in cultured iPSC-CMs. **a**) Timeline of the iPSC to CMs differentiation and proliferation experiments. **b**) Representative image of the iPSC-CMs grown with CHIR990221, staining: DAPI (blue), pHH3 (red), EdU (green), and α Actinin (yellow). Scale bar = 50 μM. **c-d**) Percentage of cells positive for proliferation markers, EdU (c) and pHH3 (d). **e-g**) Distribution of genomic content and corresponding cell cycle stages across EdU+ (f) and EdU- (g) cells based on DAPI intensity (A.U.). **e**) Distribution of genomic content according to drug concentration and EdU signal. **f**) Frequency of cell phase in EdU+ cells (2000 cells/replicate, n = 3 replicates, mean ± SD). A two-way ANOVA with Tukey’s post hoc test was used to compare the percentages of cells with a given genomic content/cell cycle stage in EdU+ cells. For G1 frequency, *P_adj_* = 0.0003 (0 μM vs. 2 μM), *P_adj_* < 0.0001 (0 μM vs. 4 μM), and *P_adj_* < 0.0001 (2 μM vs. 4 μM). For S phase, *P_adj_* = 0.0001 (0 μM vs. 4 μM) and *P_adj_* = 0.0055 (2 μM vs. 4 μM). For G2/M frequency, *P_adj_* = 0.0003 (0 μM vs. 2 μM), *P_adj_* < 0.0001 (0 μM vs. 4 μM), and *P_adj_* = 0.0003 (2 μM vs. 4 μM). For >4n frequency, *P_adj_* = 0.2685 (0 μM vs. 2 μM), *P_adj_* = 0.3236 (0 μM vs. 4 μM), and *P_adj_* = 0.9916 (2 μM vs. 4 μM). **g**) Frequency of cell phase in EdU-cells (2000 cells/replicate, n = 3 replicates, mean ± SD). A two-way ANOVA with Tukey’s post hoc test was used to compare the percentages of cells with a given genomic content/cell cycle stage in EdU+ cells. For G1 frequency, *P_adj_* < 0.0001 (0 μM vs. 2 μM) and *P_adj_* < 0.0001 (0 μM vs. 4 μM). For S phase, *P_adj_* = 0.0009 (0 μM vs. 2 μM), *P_adj_* < 0.0001 (0 μM vs. 4 μM) and *P_adj_* = 0.0291 (2 μM vs. 4 μM). For G2/M or >4n frequency, no significant differences were measured. **h**) Median DAPI intensity (A.U., a marker for cell cycle and ploidy) was measured across all replicates (2000 cells/replicate, n = 3 replicates, mean ± SD). A one-way ANOVA with Tukey’s post hoc test was used to compare the percentages of EdU+, pHH3+ cells, and median DAPI intensity. *P_adj_* is displayed in the figure. See also **Supplemental Table S1**.

We also observed significant changes in genomic content and cell cycle distribution across cells that were actively proliferating (EdU+) or non-proliferative (EdU-) during the last 48 hours of culture (**Fig. 1e-g**). Measurement of total nuclear DNA content based on total DAPI intensity is frequently used as a proxy to characterize ploidy and associated stages of the cell cycle (Pereira et al. 2017). Cardiac cell ploidy increases with development and age (Mollova et al. 2013). We saw a significant reduction in 4N (G2) frequency and an increase in 2N (G1) frequency with CHIR treatment (**Fig. 1f)**. G2 frequency decreased in EdU+ cells from 67.8% (0 µM) to 50.1% (2 µM) and 32.3% (4 µM). G1 frequency increased in EdU+ cells from 12.0% (0 µM) to 29.8% (2 µM) and 34.0% (4 µM). Alteration in genomic content and cell cycle stage could be observed independent of EdU incorporation, resulting in slight decreases in median genomic content (**Fig. 1g-h**). Examples of mononucleated and binucleated cardiomyocytes were observed at all doses of CHIR99021. We concluded that our approach provides a robust and reproducible model to generate immature CMs to study associated gene regulatory mechanisms.

### Integrated analysis of ATAC-sequencing and RNA-sequencing identifies a putative cardiac enhancer

Using the model of GSK inhibition (GSK3i), we grew differentiated iPSC-CMs with 0 µM or 4 µM CHIR for 1 week and performed the Assay for Transposase Accessible Chromatin with high-throughput sequencing (ATAC-seq) and RNA sequencing (RNA-seq). ATAC-seq measures chromatin accessibility as a marker for active or poised CREs (Grandi et al. 2022). Combining this information with RNA-seq can identify differentially accessible candidate CREs (cCREs) proximal to differentially expressed genes that respond to cell stimuli. We observed reproducible signals across all replicates in the ATAC-seq and RNA-seq data (**Supplemental Fig. S1a**). In response to GSK3i, there were 976 and 851 ATAC-seq peaks with increased or decreased accessibility, respectively, and 1023 and 1658 genes with increased or decreased expression, respectively (*P_adj_* < 0.01 and |log_2_(Fold-Change)| > 1) (**Fig. 2a-b, Supplemental Fig. S1b, Supplemental Table S2**).

**Figure 2.**
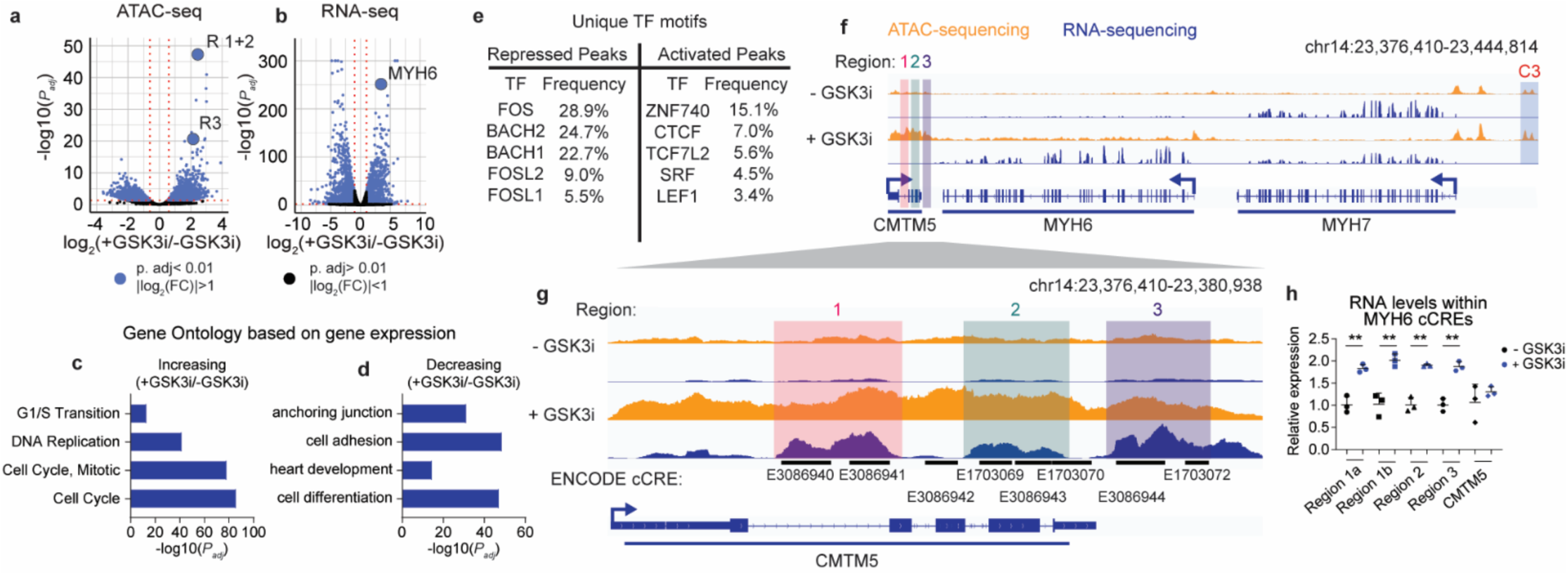
Integrated analysis of ATAC-sequencing and RNA-sequencing identifies a putative cardiac enhancer. **a-b**) Significance (*P_adj_*) versus log_2_(FC) in chromatin accessibility (a) or gene expression (b). Differential analysis was performed using a paired two-tailed DESeq2 test with a negative binomial generalized linear model and Wald statistics. **c-d**) Gene Ontology enrichment analysis for differentially expressed genes increasing (c) or decreasing (d) in expression. All Gene Ontology pathways were considered, and select key terms are displayed. **e**) The top five most frequent unique TF motifs for repressed or activated ATAC peaks with *P_adj_*<0.05. **f**) Visualization of RNA-seq (blue) and ATAC-seq (orange) results in the genomic region surrounding the *MYH6* gene body. Highlighted are three potential cCRE regions: 1 (red), 2 (green), and 3 (purple), and a previously characterized *MYH7* enhancer C3 (blue). **g**) Visualization of RNA-seq (blue) and ATAC-seq (orange) results for two significant differentially accessible peaks. **h**) RT-qPCR analysis of transcript abundance for RNAs overlapping cCRE regions and coding CMTM5 transcript levels in iPSC-CMs grown with or without GSK3i. Relative expression is plotted (normalized to TBP and -GSK3i, n=3 replicates, mean ± SD). Statistics were calculated on dCt values (normalized to TBP), using a one-way ANOVA with Sidak’s post hoc test to compare gene expression: *P_adj_*=0.0046 (region R2), *P_adj_*=0.0011 (region R1), *P_adj_*=0.0026 (region R1), *P_adj_*=0.0032 (region R1), and *P_adj_*=0.4355 (CMTM5). See **Supplemental Fig. S1** and **Supplemental Table S2**.

We further investigated the CHIR-associated changes in chromatin accessibility and gene expression. Gene ontology (GO) terms associated with increased genes included proliferation, and terms associated with decreased genes included factors associated with maturation and cardiac function, suggesting that the GSK3i-treated iPSC-CMs were in an immature state (**Fig. 2c-d, Supplemental Table S2**). Next, we investigated transcription factors (TFs) that may drive enhancer activation and alter gene expression. We applied thresholds for false discovery rate on TF motifs within peaks (FDR < 0.1) and expression (transcripts per million (TPM) > 1) for TF binding motif enrichment within ATAC-seq peaks (**Supplemental Fig. S1c-d, Supplemental Table S2**) (Bailey et al. 2015a; Fornes et al. 2020). We found that motifs corresponding to AP-1 TFs in the FOS and JUN families along with Wnt inhibitors BACH1 and BACH2 were enriched in peaks with decreasing accessibility but not in peaks with increasing accessibility following GSK3i (**Fig. 2e**). Recent reports found that these factors play roles in cardiomyocyte maturation (Zhang et al. 2023). Therefore loss of accessibility in these peaks is consistent with our model system of inducing an immature phenotype in iPSC-CMs. Motifs for TCF/LEF-associated Wnt response factors and serum response factors (SRF) were enriched in peaks increasing in accessibility (**Fig. 2e**). These factors are crucial for cardiac development and are expressed in immature CMs (Deshpande et al. 2022; Ye et al. 2019). Notably, CTCF binding motifs were enriched in peaks increasing in accessibility in response to GSK3i. CTCF, in complex with cohesin, helps determine the 3D chromatin conformation of genomic DNA. CTCF is thought to function as an insulator, preventing unintended interactions of distal enhancers with gene promoters (Cavalheiro et al. 2021). The loss of developmentally regulated CTCF binding has been observed during lineage commitment, and cohesin plays a vital role in muscle-specific gene expression (Tsai et al. 2018; Beagan et al. 2017). Additionally, CTCF can mediate enhancer-promoter interactions and gene expression outside of its canonical role as an insulator element (Kubo et al. 2021).

In response to GSK3i, we observed a shift in myosin expression from *MYH7* to *MYH6*; the myosin isoform expressed more highly in fetal cardiomyocytes (Gacita et al. 2021). *MYH6* was the 3^rd^ most significant gene to increase in expression with GSK3i with log_2_(FC) = 3.41 (*P_adj_*= 1.06×10^-251^) and *MYH7* had a log_2_(FC) = -1.340244 (*P_adj_*= 0.0036). The most significant differentially accessible peak (*P_adj_* = 6.65×10^-48^) and the 8^th^ ranked peak (*P_adj_* = 1.94×10^-48^) are located downstream of *MYH6* and *MYH7* (**Fig. 2f**). We have marked these as regions R1 through R3. We also found a previously characterized *MYH7* enhancer, which we labelled C3, to increase in accessibility following GSK3i (ranked 786^th^, *P_adj_* = 0.0007) (Gacita et al. 2021). Interestingly, we found increased RNA-seq reads overlapping R1-R3 that are not overlapping exons but rather are located within intronic and intergenic regions (**Fig. 2g**). These signals overlap cCREs across various cell types from the ENCODE database (**Supplemental Fig. S2a, c**) (Abascal et al. 2020). By RT-qPCR, we found a significant increase in these RNAs in response to GSK3i, whereas there was no change in levels of spliced coding transcripts for the overlapping gene *CMTM5* (**Fig. 2h**). Transcription of noncoding regulatory elements is a marker of active enhancers and can help promote enhancer-promoter interactions (Sartorelli and Lauberth 2020; Tsai et al. 2018). In the VISTA enhancer browser, this region shows no activity during mouse development, throughout which *Myh7* is the dominant myosin isoform (Visel et al. 2007; May et al. 2012).

### Epigenetic landscape surrounding a gene regulatory element in human and mouse samples

To understand the context under which this putative regulatory element may be active, we analyzed multiple heart epigenetic data sets from human and mouse cardiac samples and non-heart data sets from humans. We find that across tissue types, neuronal tissues, followed by heart tissue types, have the highest H3K27ac signal in regions R1-R3 (**Supplemental Fig. S2a**). The high activity in neuronal tissue types is likely driven by the high expression of CMTM5 (**Supplemental Fig. S2b**).

The genomic composition surrounding *MYH6* and *MYH7* is preserved between humans and mice (**Supplemental Fig. S3a**). The genes and CTCF binding sites within this region are the same, with identical orientations. In both human and mouse heart samples, there is a CTCF ChIP-seq signal within R3 as well as a distal CTCF-bound enhancer (labeled M3), and another peak is proximal to the previously characterized C3 myosin enhancer (**Fig. 3a, Supplemental Fig. S3a**). The C3 enhancer is essential for *MYH7* expression and has a minor role in *MYH6* expression (Gacita et al. 2021). Single-cell ATAC-seq from atrial or ventricular cardiomyocytes also shows similarities to our iPSC-CM ATAC-seq peaks (**Fig. 3a**) (Hocker et al. 2021a). In both atrial and ventricular cardiomyocytes, we observe chromatin accessibility around R1-R3. Next, we looked at histone marks associated with enhancer activity. H3K27me3, a marker of enhancer repression, was present in R3 in the ventricular samples but depleted in the atrial samples (**Fig. 3b**). We observed a signal for H3K27ac, a marker for enhancer activity, in cardiac samples overlapping R3 and throughout the gene body of *MYH6* (**Fig. 3a, Supplemental Fig. S2a**).

**Figure 3.**
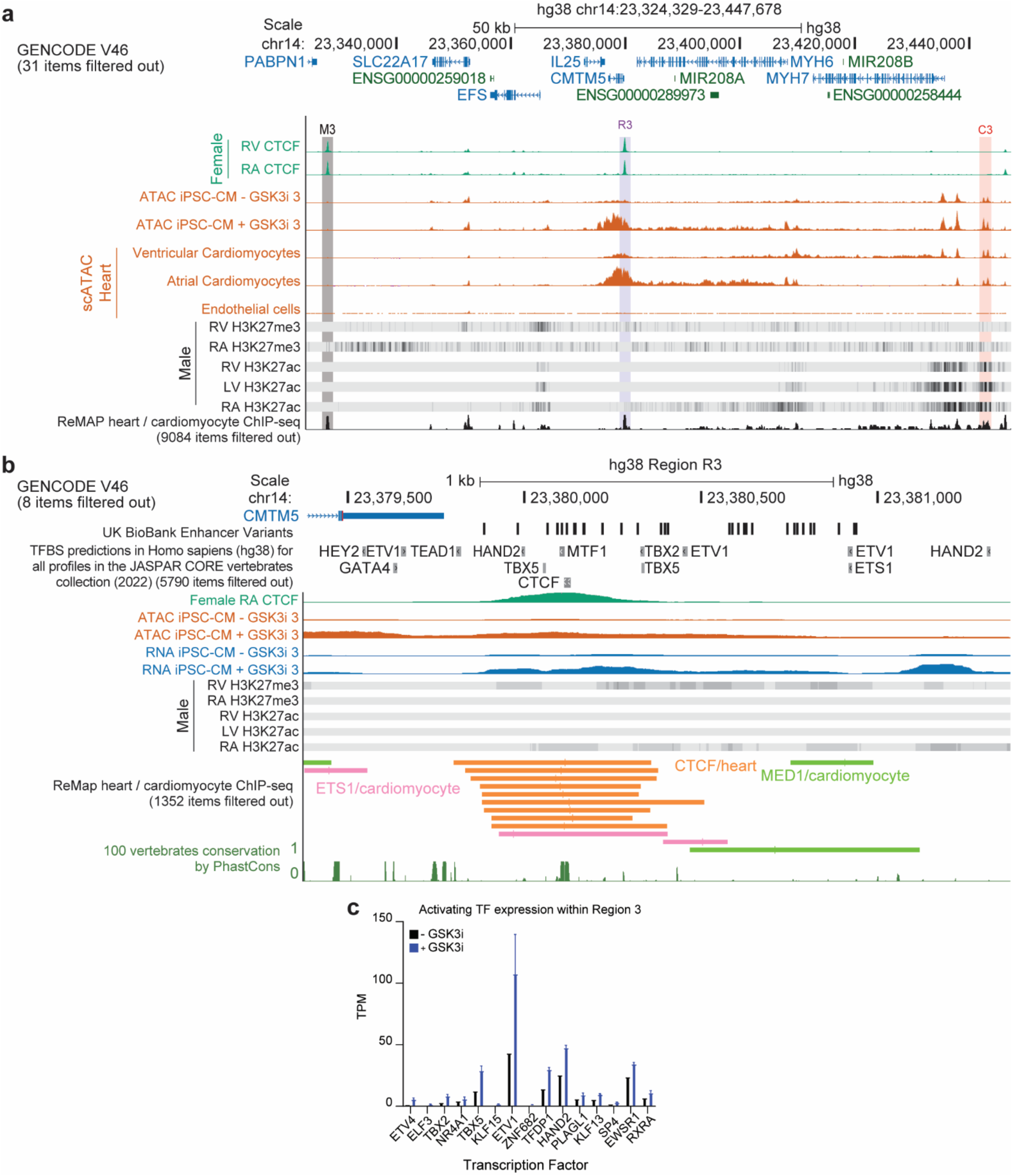
Epigenetic landscape surrounding a cardiac-specific candidate *cis*-regulatory element (cCRE) in human samples. **a-b**) Visualization of epigenetic datasets surrounding the *MYH6* locus (a) and region R3 (b). Datasets include CTCF-ChIP (green), ATAC-seq profiles from iPSC-CMs and single-cell ATAC-seq in human hearts (orange), RNA-seq profiles in iPSC-CMs (blue), histone ChIP-seq density plots, and ReMap ChIP-seq data from cardiac-related studies. The cCRE regions are highlighted as follows: M3 (grey), R3 (purple), and C3 (orange). **c**) TPM values for transcription factors (TFs) with binding sites within R3 (FDR < 0.01) that significantly increase in expression following GSK3i treatment (*P_adj_*< 0.01 and log_2_(FC) > 1). See also **Supplemental Fig. S2, S3, S4**.

Within R3 specifically, we find TF motifs for factors necessary for cardiac gene expression (TBX5, TBX2, HAND2, GATA4, HEY2, ETV1, ETS1) (Ang et al. 2016; Tsuchihashi et al. 2011; Rommel et al. 2018; Shekhar et al. 2016). Based on cardiac ChIP data from the ReMAP Atlas, ETS1 and MED1 bind R3 in cardiomyocytes (**Fig. 3b**) (Hammal et al. 2022). TBX5, GATA4, and MED1 can form a complex important for cardiac homeostasis (Ang et al. 2016). In mice, R3 contains similar TF motifs and is bound by TFs p300 and Tead1 based on ChIP-seq in adult mouse hearts (**Supplemental Fig. S3b**) (Ghosh 2020). p300 disruption is embryonically lethal and linked to defects in heart development (Shikama et al. 2003). Activation of p300 occurs during heart failure, and its overexpression leads to increases in Myh7 expression in mice (Wei et al. 2008). The histone and TF ChIP data suggest that R3 likely functions as an enhancer and has activity in the heart and neuronal cells.

The previously characterized enhancer, C3, and the CTCF-bound M3 enhancer both harbor distinct binding motifs. The C3 enhancer is bound by TFs that are important for cardiac development (GATA4, TBX5, MED1, and ETS1) (**Supplemental Fig. S4a, c**). Both C3 and M3 contain binding motifs for BACH2, as well as MEF2C, which has been shown to be an important TF for cardiac function (**Supplemental Fig. S4b, d**) (Vincentz et al. 2008). Recent evidence suggests that CTCF is essential for gene expression and promoter-enhancer interactions for a subset of genes (Kubo et al. 2021). We assessed the expression of the TFs with motifs in R3 in our RNA-seq data and observed increased expression of *ETV1*, *HAND2*, *TFDP1*, *EWSR1*, and *TBX5* (**Fig. 3c**). The activation of these TFs may be involved in the increased activity of R3. TBX5 is crucial for *MYH6* expression (Ching et al. 2005). However, the role of these other TFs in *MYH6* expression has not been directly investigated.

We analyzed publicly available Hi-C data to investigate *MYH6* promoter interactions in cardiac samples, which we obtained from the 3D-genome Interaction Viewer and Database (3DIV) (Jung et al. 2019; Yang et al. 2018). We found that the *MYH6* promoter interacts with all three described enhancers, C3, R1-R3, and M3 (**Supplemental Fig. S5**). The *MYH6* promoter exhibits more local interactions looping to the C3 enhancer region preferentially in left ventricular CMs and loops to the distal M3 enhancer in right ventricular CMs (**Supplemental Fig. S5a-b**). We also observed that R1-R3 interacts with the *MYH6* promoter (**Supplemental Fig. S5c-d**).

Based on the analysis of single-cell chromatin accessibility, histone marks, chromatin looping, TF ChIP-seq, and motif analysis, R3 and M3 have profiles similar to the previously characterized *MYH7* enhancer, C3, and physically interact with *MYH6*. This data provides evidence that these regions may work in concert to regulate *MYH6* and *MYH7*.

### CRISPR-mediated epigenetic editing of R3 modulates cellular response to disease-related stimuli

Given the genomic indicators of cardiac activity of R1-R3, we next sought to link the activity of this region to the expression of its target gene. To achieve this, we epigenetically repressed this enhancer with dCas9^KRAB^ (CRISPRi) or activated it with ^VP64^dCas9^VP64^ (CRISPRa) and analyzed gene expression by RT-qPCR (**Fig. 4a**). These CRISPR epigenome editors can effectively alter gene expression when delivered to the enhancer of a target gene (McCutcheon et al. 2023a). We transduced iPSC-CMs with lentivirus expressing dCas9^KRAB^ and a gRNA targeting the enhancer of interest or a control gRNA with no target in the human genome. At twelve days following transduction, we found that repression of regions R1 and R3 decreased *MYH6* expression to 76% and 49% of controls, respectively, while region R2 did not affect expression (**Fig. 4b**). Repression of the enhancers had no impact on *MYH7* expression (**Fig. 4c**). A marker of cardiomyocyte maturation is a shift in expression from *MYH6* to *MYH7* (Gacita et al. 2021). Enhancer repression shifted the *MYH7/MYH6* ratio from 2.25 in control samples to 3.02 (R1), 2.49 (R2), and 4.29 (R3) when the enhancer regions were targeted. CMTM5 levels dropped to 60% and 73% of controls when regions 2 and 3 were targeted, respectively (**Fig. 4d-e**). This links R3 activity to MYH6 expression and shows that MYH6 expression is not regulated by CMTM5, as repression of region R2 decreased CMTM5 expression but not MYH6 expression. We observed a slower rate of spontaneous contraction following repression of R3 (**Supplemental Video 1** = control, **Supplemental Video 2** = R3 repression). It has been previously reported that alterations in myosin isoform abundance affect the rate of contractions of iPSC-CMs (Gacita et al. 2021). The remaining experiments focused on R3 due to the more significant observed effect of the perturbation of R3 on myosin expression.

**Figure 4.**
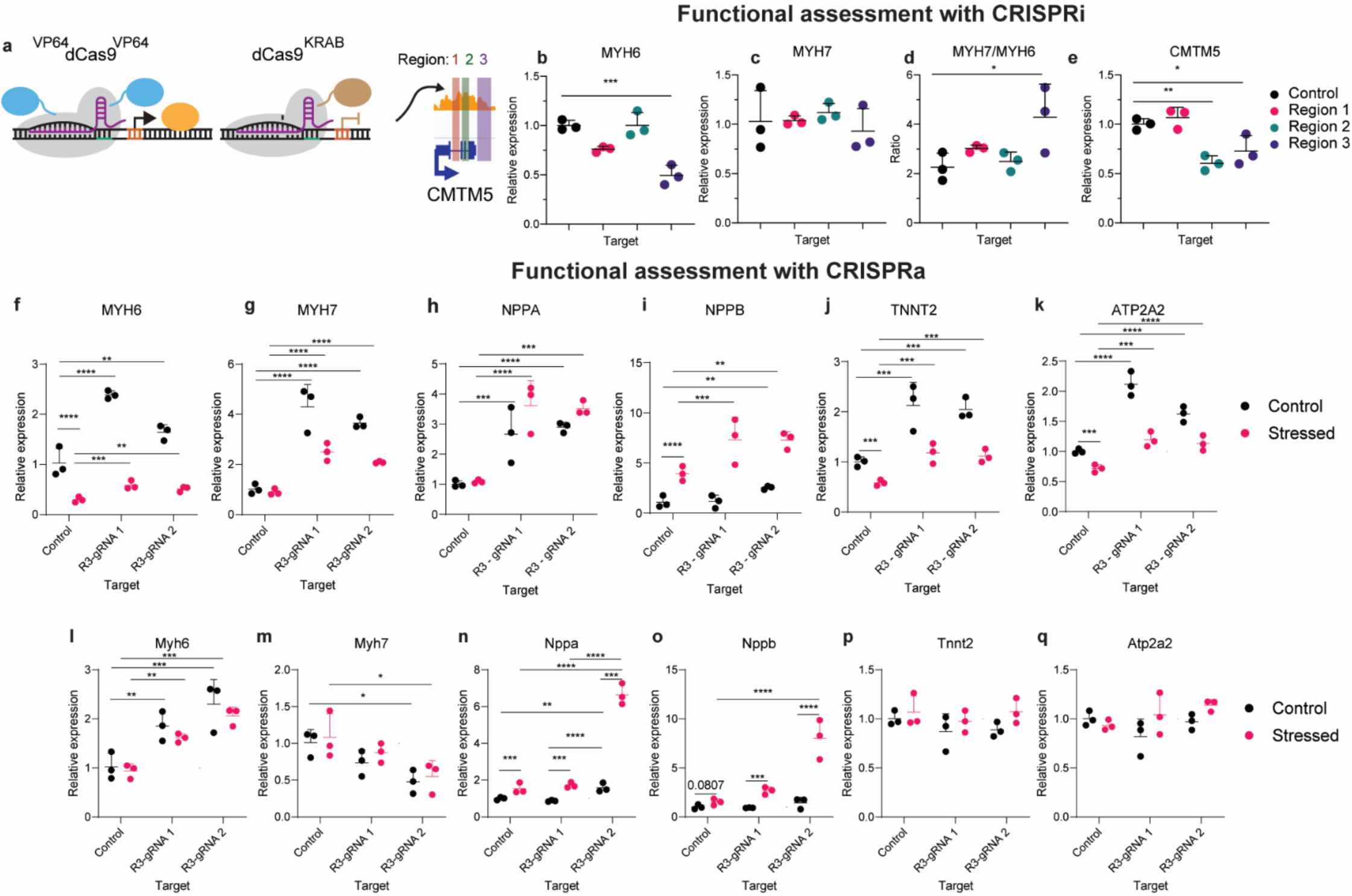
CRISPR-mediated epigenetic editing of R3 alters cellular response to disease-related stimuli. **a**) Schematic of cCRE targeting using CRISPR epigenome editors **b-e**) RT-qPCR for MYH6 (b), MYH7(c), MYH7/MYH6 (d), and CMTM5 (e) 12 days after repression of each designated cCRE. Relative expression is plotted (normalized to TBP and control, n = 3 replicates, mean ± SD). Statistics were calculated on dCt values (normalized to TBP). A one-way ANOVA with Dunnett’s post hoc test was used to compare gene expression: *P_adj_* =0.0003 (MYH6, control vs region R3), *P_adj_*=0.0239 (MYH7/MYH6, control vs region R3), *P_adj_*=0.0045 (CMTM5, control vs region R3), and *P_adj_*=0.0404 (CMTM5, control vs region R3), **f-k**) RT-qPCR for MYH6 (f), MYH7(g), NPPA (h), NPPB (i), TNNT2 (j), and ATP2A2 (k) 15 days following activation of each designated cCRE in human iPSC-CMs ± ET-1 1 μM for 72 hours. Relative expression is plotted (normalized to TBP and control, n = 3 replicates, mean ± SD). Statistics were calculated on dCt values (normalized to TBP), and a two-way ANOVA with Tukey’s post hoc test was used to compare gene expression. **l-q**) RT-qPCR for Myh6 (l), Myh7(m), Nppa (n), Nppb (o), Tnnt2 (p), and Atp2a2 (q) 15 days following activation of each designated cCRE in HL-1 mouse atrial cardiomyocytes ± Norepinephrine and FBS for 48 hours. Relative expression is plotted (normalized to TBP and control, n = 3 replicates, mean ± SD). Statistics were calculated on dCt values (normalized to TBP), and a two-way ANOVA with Tukey’s post hoc test was used to compare gene expression.

Next, we determined the effects of enhancer activation on iPSC-CMs under normal and stressed culture conditions. *MYH6* expression is important for decreases in heart function during heart failure and under stress (Ching et al. 2005; Chen et al. 2021; Han et al. 2014). We chose endothelin-1 (ET-1) treatment to model cardiac stress. ET-1 is a GPCR agonist and vasoconstrictor released by endothelial cells to drive cardiac hypertrophy (Dhaun et al. 2008; Mueller et al. 2011). ET-1 also represses *MYH6* and reactivates fetal gene expression profiles that occur during heart failure (Bloch et al. 2016; Jiang et al. 2019; Man et al. 2021; Talman and Kivelä 2018). We transduced iPSC-CMs with a lentiviral vector encoding the activator ^VP64^dCas9^VP64^ and a gRNA targeting R3. Twelve days after transduction, the cells were treated with 1 µM ET-1 or vehicle (H_2_O) for 72 hours. Activation of R3 significantly increased expression of *MYH6* and *MYH7* compared to control gRNA across culture conditions with two independent gRNAs; ET-1 treatment diminished the activation of both genes, but R3-activated cells maintained higher expression of these genes than the cells with the control gRNA (**Fig. 4f-g**).

*NPPA* and *NPPB* are essential factors during cardiac development and are markers of stress and pathologic hypertrophy (Kerkelä et al. 2015b; Man et al. 2021). These two factors reactivate in response to stress or signaling molecules such as ET-1 (Sergeeva et al. 2016a; Jiang et al. 2019). To understand how the shifts in MYH6/7 expression with and without stress affect these downstream markers, we also analyzed NPPA and NPPB expression. NPPA expression increased with myosin activation independent of culture conditions (**Fig. 4h**). NPPB increased only in the presence of ET-1 and had a more significant increase in R3-activated cells (**Fig. 4i**) (Jiang et al. 2019). The increases in expression of NPPB under stress indicate that R3 activation may sensitize iPSC-CMs to ET-1 stress.

Increased expression of *TNNT2* and *ATP2A2*, which are on different chromosomes from *MYH6* and *MYH7*, occurs with increased myosin expression and is maintained following ET-1 exposure (**Fig. 4j-k**). *TNNT2* and *ATP2A2* encode proteins that are crucial for cardiac function. *TNNT2* encodes cardiac troponin T, and this factor is mutated in a subset of inherited cardiomyopathies (Crocini and Gotthardt; Parbhudayal et al. 2020). *ATP2A2* encodes SERCA2a, which is vital for calcium handling in the heart, and its disruption occurs with many forms of cardiac disease (Gilbert et al. 2020; Kumarswamy et al. 2012). ET-1 reduces the expression of SERCA2a, which results in decreased cardiomyocyte function and prolonged calcium signaling, similar to heart failure (Uehara et al. 2012; Mueller et al. 2011). We observed that R3 activation results in higher expression of both *TNNT2* and *ATP2A2* than cells with the control gRNA in control and stressed culture conditions.

To evaluate if this regulatory mechanism is conserved across species, we evaluated enhancer function in the mouse HL-1 atrial cardiomyocyte cell line that is often used to study heart disease (Bloch et al. 2016; Claycomb et al. 1998). We used norepinephrine and FBS to induce a hypertrophic phenotype, which resulted in a similar increase in *NPPA* expression and a more significant increase in *NPPB* than ET-1 (**Supplemental Fig. S6c-d**) (Bloch et al. 2016; Dambrot et al. 2014). Enhancer activation by CRISPRa resulted in expression changes that were similar to those of human iPSC-CMs. We observed changes in *Myh6*, *Nppa*, and *Nppb* that are consistent with those in the human context (**Fig. 4l, n-o**). However, we observed decreased *Myh7* expression (**Fig. 4m**). This may be related to underlying differences in biology between humans and mice: *Myh6* is the dominant isoform in adult mice, and *MYH7* is the dominant isoform in adult humans (Miyata et al. 2000). This difference may result from divergent mechanisms of myosin regulation and gene expression that have evolved over time. Finally, we observed no changes in *Tnnt2* or *Atp2a2* expression following enhancer activation in HL-1 cells (**Fig. 4p-q**) (Claycomb et al. 1998).

Based on the above data, R3 activity is important for myosin expression. R3 repression decreased *MYH6* expression, and activation led to increased expression of *MYH6* and *MYH7*. The increased expression of myosin through R3 activation leads to activation of other cardiac genes and an altered cellular response to stress caused by ET-1. These key cardiac factors are related to sarcomere composition, calcium signaling, and paracrine signaling, which may alter cardiomyocyte composition and function.

### Activation of R3 alters cellular composition and function in control and stressed states

Having linked R3 activity to myosin expression and finding that this perturbation results in altered expression of multiple cardiac-specific factors, we sought to quantify phenotypic effects driven by R3 activation in control and stressed culture conditions. ET-1 can influence cell size and electrophysiology in culture (Bourque et al. 2022; Watkins et al. 2011; Uehara et al. 2012). Cardiac polyploidization, marked by increases in genomic content, is linked to both physiological and disease-related cardiac hypertrophy (Gilsbach et al. 2018; Derks and Bergmann 2020; Ya Brodsky -Donat Sarldsov et al. 1994; Bergmann et al. 2015).

We measured changes in genomic content and cell size in human iPSC-CMs following R3 activation in control and stressed culture conditions (**Fig. 5a**). We found that R3 activation did not significantly alter genomic content under control conditions (*P*=0.382) but found that it significantly reduced the increase in genomic content that occurred following stress with ET-1 for 48 hours (*P*=0.0026) (**Fig. 5b**). Cell size did not change substantially following R3 activation (**Supplemental Fig. S6a**). Culture with ET-1 increased cell size across control and R3 gRNA conditions and R3-activated cells were slightly smaller than control counterparts (*P*=0.3523) (**Supplemental Fig. S6a**). We found that when we performed linear regression on cell size and total genomic content of each cell, the slope was greater for cells following R3 activation across culture conditions, implying that for equal genomic content, R3-activated cells should generally be larger (**Supplemental Fig. S6b**). When cell size was normalized by genomic content, R3-activated cells are significantly larger under stressed conditions (*P*=0.0254) and have trending differences in control conditions (*P*=0.0891) (**Fig. 5c**). We looked at the relative abundance of MYH7 to MYH6 (a marker of iPSC-CM maturity) and found that R3 activation increased expression of both myosin isoforms and shifted the relative abundance toward MYH7, suggesting increased maturity (**Fig. 4f-g**, **Fig. 5d**) (Gacita et al. 2021). We also assessed the difference in cell size of HL-1 cells by flow cytometry under stressed conditions. We performed flow cytometry on HL-1 cells 3 weeks following enhancer activation (**Supplemental Fig. S6c-e**). We observed similar differences in the HL-1 cells as the human iPSC-CMs, with up to a 16.3% decrease in median cell size by forward scatter following enhancer activation (*P_adj_=0.0714)* (**Fig. 5e**).

**Figure 5.**
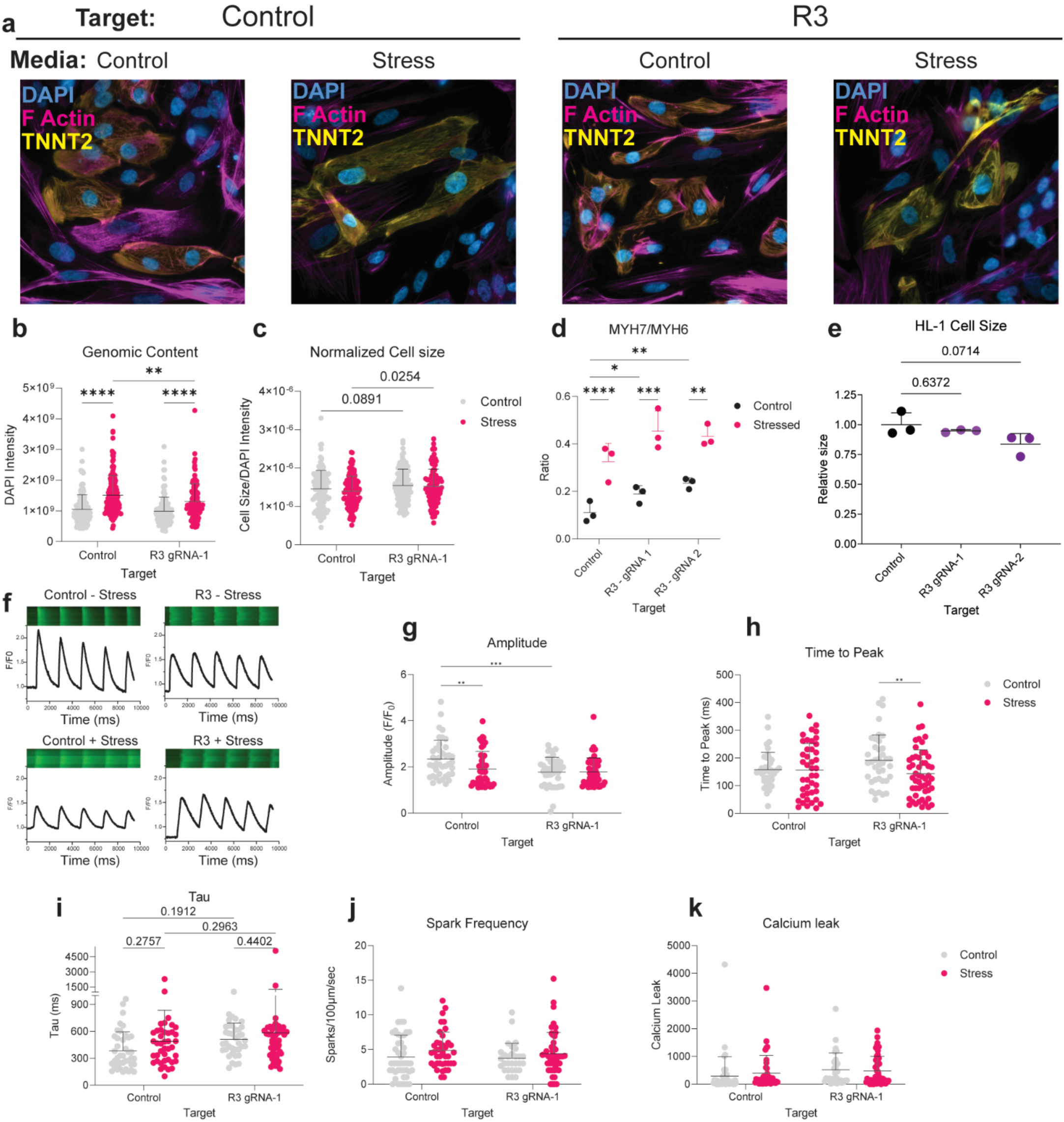
Activation of R3 alters cellular composition and function in control and stress states. **a**) Representative images of iPSC-CMs receiving control or R3 gRNA-1, grown under control or stressed conditions. **b-c**) Analysis of genomic content (b) and cell size (c) of human iPSC-CMs represented in (**a**) 20 days following activation of each designated target ± ET-1 (1 μM) for the last 48 hours of culture, n = 3 replicates. Statistics were calculated with a two-way ANOVA with Fisher’s LSD. **b**) Measurement of genomic content (DAPI intensity within the area of the cell). **c**) Cell size normalized by the genomic content of each cell. **d**) The ratio of MYH7 to MYH6 abundance was measured using the data from **Fig. 4f-g**. Ratios are plotted (normalized to TBP and control, n = 3 replicates, mean ± SD). Statistics were calculated on dCt values (normalized to TBP), and a two-way ANOVA with Tukey’s post hoc test was used to compare ratios. **e**) Relative cell size measured 21 days following activation of each designated cCRE in HL-1 mouse atrial cardiomyocytes ± Norepinephrine + FBS (median FSC-A normalized to the control, n = 3 replicates, mean ± SD). A one-way ANOVA test with Dunnett’s post hoc test was used to compare cell size: *P_adj_* = 0.6372 (Control vs. R3 gRNA-1) and *P_adj_* = 0.0714 (Control vs. R3 gRNA-2). **f**) Representative confocal line scan image and corresponding Ca^2+^ fluorescent tracing in iPSC-CM receiving control gRNA or R3 gRNA grown under control or stressed conditions for 24 hours. **g-k**) Dot plots for quantitative analysis of cells represented in (**f**) (n = 6 replicates across two differentiations). Statistics were calculated with a two-way ANOVA with Fisher’s LSD. **g**) Dot plots for Amplitude (F/F_0_). **h**) Dot plots for time to peak. **i**) Dot plots for Tau. **j**) Dot plots for spark frequency. **k**) Dot plots for calcium leak. (\**P_adj_* < 0.05, \*\**P_adj_* < 0.01, \*\*\**P_adj_* < 0.001, \*\*\*\**P_adj_* < 0.0001). See also **Supplemental Fig. S6**.

ET-1 has been shown to drive alterations in electrophysiology that precede the occurrence of heart failure in mouse models (Mueller et al. 2011). We also observed differences in calcium handling protein ATP2A2 expression levels following R3 activation across culture conditions (**Fig. 4k**). Therefore, we sought to measure electrophysiological differences between iPSC-CMs ± R3 activation across culture conditions. We measured intracellular Ca^2+^ dynamics in iPSC-CMs using the fluorescent dye Cal-520 (**Fig. 5f**) (Daily et al. 2017). We found that the spontaneous Ca2+ transient (F/F_0_) in iPSC-CMs was reduced following R3 activation (*P*=0.0004) (**Fig. 5g**). The amplitude (F/F_0_) decreased significantly following stress with ET-1 in cells with the control gRNA (*P*=0.0057) but did not decrease for R3-activated cells following stress with ET-1 (**Fig. 5g**). Time to peak shortened for R3-activated cells following stress (P=0.0086), whereas it was unchanged for control cells. (**Fig. 5h**). We observed no statistical difference between control and R3-activated cells across culture conditions for other measurements related to calcium dynamics (Tau, spark frequency, or calcium leak) (**Fig. 5i-k**). In conclusion, R3 activation alters cellular response to stress, and activated cells are functionally distinct from control cells. R3 activation attenuates polyploidization caused by ET-1 and increases normalized cell size. Ratios of MYH7 to MYH6 shift toward MYH7 as a marker of increased maturity following R3 activation (Gacita et al. 2021). R3 activation reduced the baseline amplitude for calcium dynamics and prevented the amplitude decline following stress with ET-1.

### Activity of R3 alters chromatin looping surrounding *MYH6* and *MYH7*

Based on the evidence that CTCF binding and chromatin looping may be important for the enhancer function of R3 (**Fig. 2e, 3a, 3d, Supplemental Fig. S5c-d**), we examined changes in chromatin conformation in iPSC-CMs by Hi-C of Accessible Regions (HiCAR) in combination with CRISPR-based epigenetic perturbation of R3 (**Supplemental Fig. S7**) (Wei et al. 2022). We found that R3 and C3 interact with the promoter of *MYH6* (**Fig. 6a-b, Supplemental Table S2**). We performed probe-enriched HiCAR sequencing on genomic DNA from iPSC-CMs with R3 repression or activation in stressed or control conditions and HL-1 cells under control culture conditions (Wei et al. 2022). Activation of R3 alone did not drive significant changes in chromatin conformation (**Fig. 6c**) therefore the increase in *MHY6* and *MYH7* expression following targeting of CRISPRa to R3 (**Fig. 4f, g**) may involve more efficient activation of *MYH6* through preexisting loops. However, the repression of R3 by CRISPRi resulted in a significant increase in interactions between the promoter and gene body of *MYH6* with the distal CTCF-bound enhancer (M3) (**Fig. 6d**). This recapitulates the chromatin conformation observed in right ventricular samples (**Supplemental Fig. S5b**) and indicates that targeting R3 with dCas9^KRAB^ promotes de novo MYH6/M3 looping, concomitant with loss of *MYH6* expression, while *MYH7* levels are unchanged (**Fig. 4b, c**).

**Figure 6.**
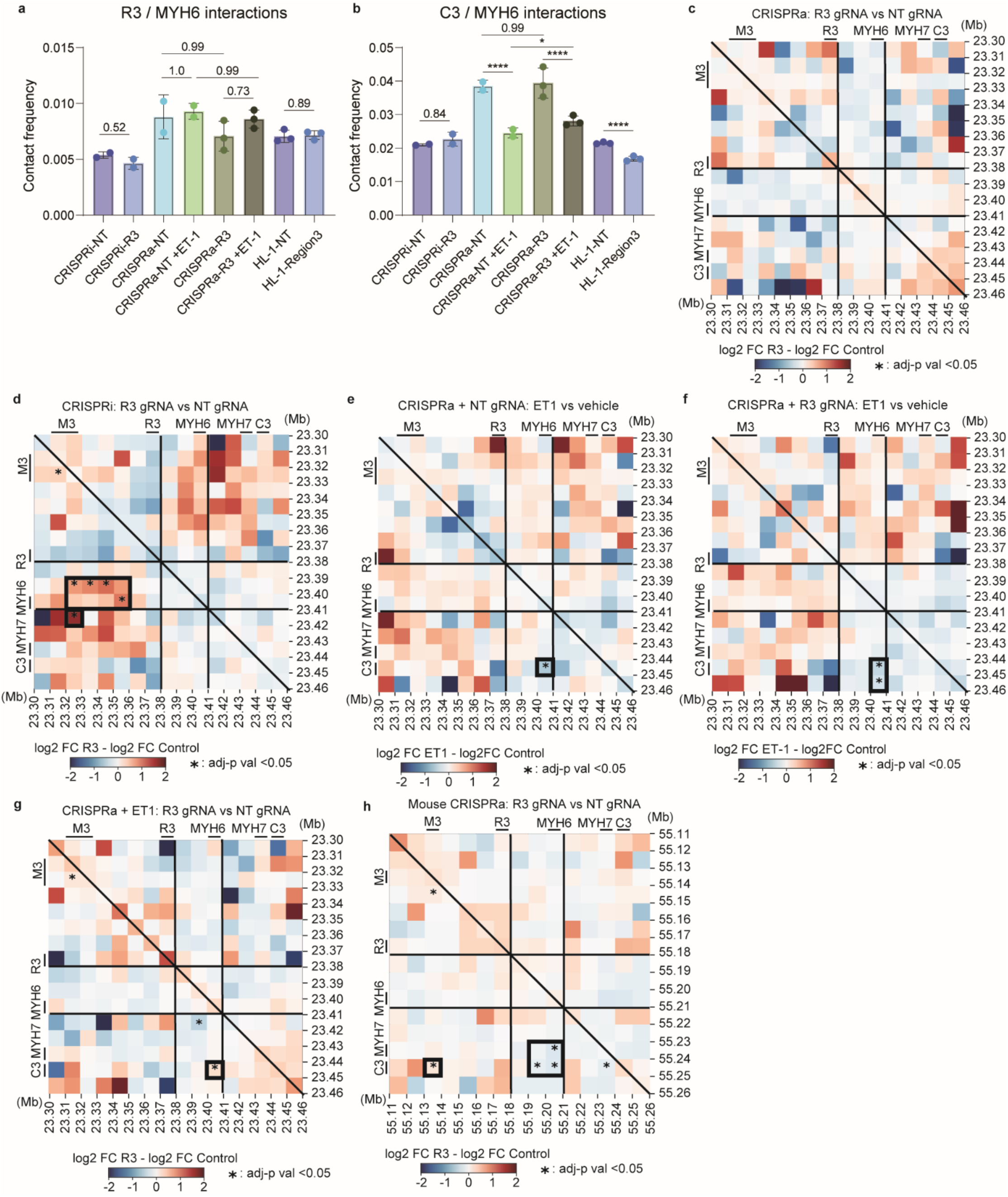
Activity of R3 alters chromatin looping surrounding *MYH6* and *MYH7*. **a-b**) Contact frequency between the MYH6 TSS and R3 (a) or C3 (b) normalized to total contacts in the region assessed during DESeq2 analysis of the contact matrix visualized in panels **c-h**. **c-h**) Differential contact frequency heatmaps showing the impact of R3 activity on chromatin looping under control and stressed conditions. Enhancer regions (R3, C3, M3) and the TSS for MYH6 and MYH7 are indicated alongside each heatmap. Differential analysis was performed using a paired, two-tailed DESeq2 test with a negative binomial generalized linear model and Wald statistics. Significant non-self-ligated contacts (*P_adj_* < 0.05) are marked with an asterisk (*). Genomic coordinates: hg38, chr14:23,303,865-23,451,365; mm39, chr14:55,111,190-55,258,690. Heatmaps compare differential contact frequency between R3 activation and control under non-stressed conditions (**c**) or R3 repression and control (**d**) in human iPSC-CMs. **e-g**) Panels display differential contact frequencies after 72 hours of ET-1 exposure: without R3 activation (**e**), with R3 activation (**f**), and a differential heatmap for R3-NT under stressed conditions (**g**). **h**) The heatmap comparing R3 activation and control in non-stressed HL-1 mouse atrial cardiomyocytes. DESeq2 results (*P_adj_* and log2FC) are provided. Regions of interest discussed in the text are highlighted with a thick border. See also **Supplemental Fig. S7** and **Supplemental Table S3**.

When we treated iPSC-CMs with ET-1, we observed significant decreases in interactions between *MYH6* and the C3 enhancer region, regardless of R3 activation, with a 38% decrease in control (*P_adj_* < 0.0001) and a 32% decrease in R3-activated cells (*P_adj_* < 0.0001) (**Fig. 6e-f**). In contrast, in the presence of ET-1, R3 activation leads to significantly more MYH6/C3 interactions than cells lacking R3 activation, with a 23% increase in total contacts (*P_adj_* = 0.023) (**Fig. 6g**). These results suggest that R3 activation directly opposes the alterations caused by ET-1 on MYH6/C3 looping and silencing of MYH6 (**Fig. 4f**).

Finally, we examined chromatin conformation within the HL-1 cells. Activation of R3 in HL-1 cells resulted in decreased interactions between the MYH6 promoter and the mouse C3 region (22% decrease, *P_adj_*

*< 0.0001)* and increased interactions between the distal M3 enhancer and C3 by 66% (*P_adj_ =0.0152)* (**Fig. 6h**). This was not observed in the human iPSC-CMs and may contribute to the underlying differences in myosin expression observed between humans and mice (Miyata et al. 2000). However, additional studies are needed to dissect the role that looping plays in the differences between mouse and human myosin expression.

We demonstrate that chamber-specific chromatin looping surrounding the *MYH6* locus can be recapitulated by altering the activity of R3 **(Fig. 6d, Supplemental Fig. S5a-b**). This supports a model in which CTCF-bound enhancer looping is vital for expression, where the activity of CTCF-bound enhancers determines promoter interactions. In the context of myosin regulation, high to moderate R3 enhancer activity maintains local interactions with the *MYH6* TSS. However, as R3 activity declines, the distal M3 enhancer interacts more frequently with the *MYH6* promoter, reducing *MYH6* expression. When iPSC-CMs reduce *MYH6* in response to stress, local interactions between *MYH6* and the C3 proximal enhancer also decline (**Fig. 7**). Forced reactivation of R3 prevents this dissociation, and cells maintain higher levels of *MYH6*.

**Figure 7.**
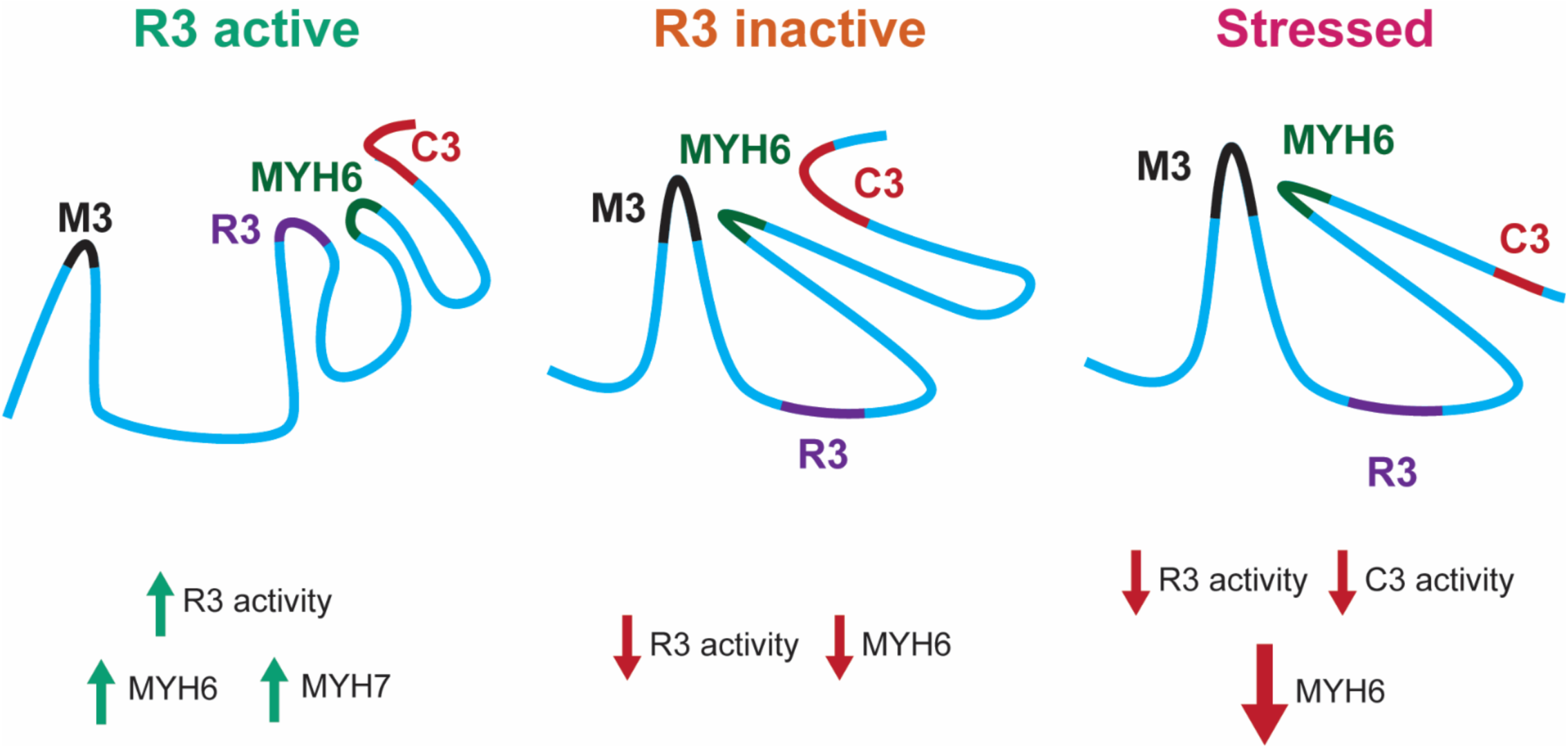
Model of Myosin Regulation. R3 activity is crucial for MYH6 expression. Varying R3 activity alters MYH6 chromatin contacts in iPSC-CMs. R3 activity results in increased MYH6 and MYH7 expression and altered functional readouts. In contrast, reduced R3 activity allows the distal M3 to interact more frequently with MYH6, decreasing MYH6 expression. Additionally, stress-induced loss of C3 contact results in additional decreases in MYH6 expression. R3 activation is antagonistic to this loss of contact, prevents further decline in calcium signaling, and reduces polyploidization resulting from stress.

### Analysis of genetic variants within the myosin enhancer

Genetic variations associated with heart disease are enriched within cardiac regulatory elements and are linked to cardiac phenotypes in model systems (Gilsbach et al. 2018; Wang et al. 2016). These variants can be hidden below previously set thresholds in genome-wide association studies (Wang et al. 2016). When this analysis is combined with epigenetic profiling and functional perturbations, new noncoding genetic variants can be linked to complex human traits. Therefore, we analyzed genetic variation within this newly characterized enhancer region in humans utilizing participant whole genome sequencing (WGS) data from the UK Biobank (**Supplemental Table S4**). We compared the frequency of variants within regions R1, R2, and R3 in patients with or without cardiomyopathies. Variations within enhancer regions can inactivate these elements, silencing target genes (Schubach et al. 2024; Jung et al. 2019). Since R3 is crucial for *MYH6* expression, and *MYH6* silencing is linked to heart failure, we hypothesized that disruptive variants within R3 may occur more frequently between participants with cardiomyopathies or control participants.

We performed a Fisher’s exact test to assess the frequency of variants within regions R1, R2, and R3 between participants with and without cardiomyopathy, excluding individuals with coronary artery disease (**Supplemental Table S4**). A significant *P_adj_* would signify that a variant occurs more frequently in either cardiomyopic or control participants. We evaluated only variants within intronic and noncoding regions overlapping R1 to R3 and plotted -log10(*P_adj_*) (frequency) against the CADD score, a measure of variant deleteriousness, to identify variants more frequent in cardiomyopathy participants with a higher likelihood of being pathogenic (**Fig. 8a**) (Schubach et al. 2024). We found that only variants within R3 had a *P_adj_* < 0.05. Next, we calculated a tolerance score for variants located in motifs of expressed TFs in R3 (**Fig. 8b**), where a negative score indicates reduced tolerance and potentially impaired TF binding, and a positive score indicates greater tolerance.

**Figure 8.**
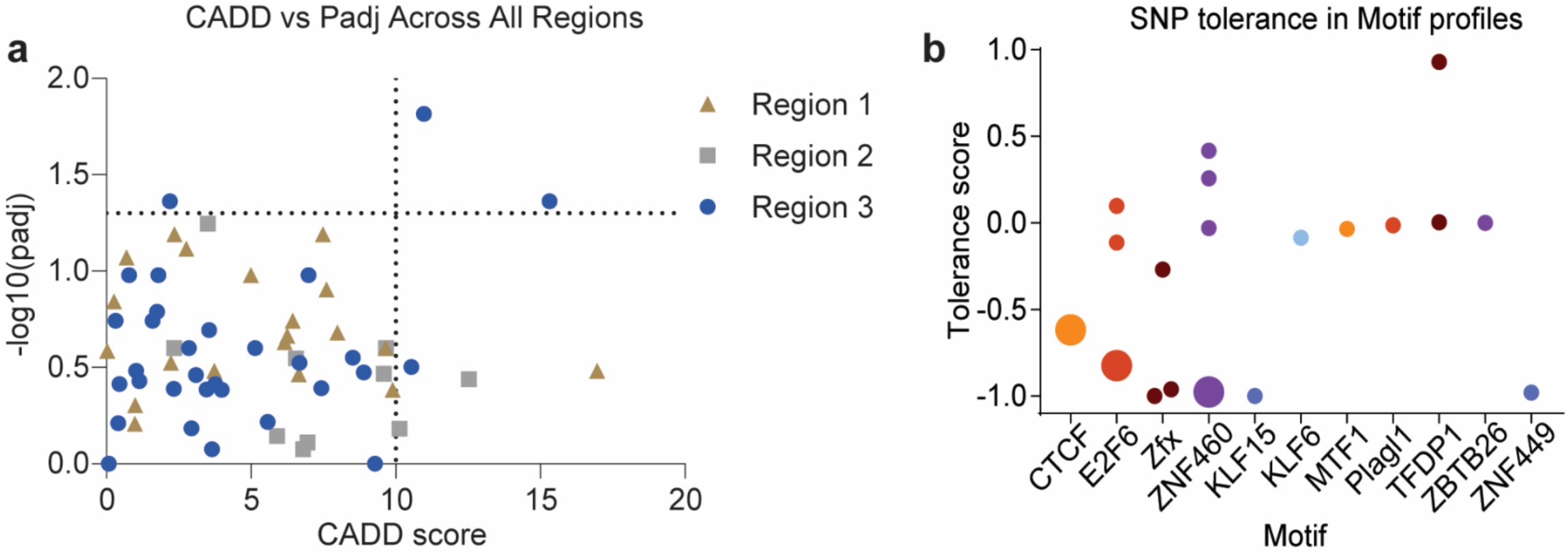
Analysis of genetic variants within the myosin enhancer. **a**) Visualization of CADD scores for variants within regions R1 - R3 found in subjects from the UKBiobank. CADD scores were determined through CADDv1.7, and the *P_adj_* was determined by a Fisher’s exact test on the frequency of genetic variants in participants with or without a cardiomyopathy. Colors and shapes correspond to the region containing the variant. The dotted lines are at *P_adj_* =0.05 and a PHRED score = 10. **b**) Tolerance scores for genetic variants within motif profiles found in R3 from human participants in the UK Biobank. Enlarged dots correspond to variants with *P_adj_* <0.05. See also **Supplemental Table S4**.

We identified a rare heterozygous variant in a highly conserved CTCF binding motif with a reduced tolerance score (-0.618) that was more common in participants with cardiomyopathy (p=0.043) and had a high CADD score (15.32), placing it in the top 10% of predicted deleterious variants (Schubach et al. 2024). Additionally, a rare homozygous variant disrupting E2F6 motifs, which showed reduced tolerance for TF binding, was enriched in cardiomyopathy participants (*P_adj_* = 0.022, CADD = 2.934). Interestingly, a heterozygous variant disrupting ZNF460 and Zfx binding was more frequent in individuals without cardiomyopathy (*P_adj_* = 0.015, CADD = 10.97).

Further studies on genetic variation in this myosin enhancer and other regulatory regions will be necessary to understand the noncoding genome’s role in heart disease and other complex human traits. Silencing of *MYH6* has also been observed to cause cardiac septal defects in chick embryo models and is associated with various cardiomyopathies (Ching et al. 2005; Lowes et al. 1997; Nakao et al. 1997; Miyata et al. 2000; Locher et al. 2011). Allelic imbalances in the expression of mutant versus wild-type *MYH7* have been observed in hypertrophic cardiomyopathy and are proposed as potential drivers of pathogenesis (Montag and Kraft; Montag et al. 2017). R3 inactivation on a single allele could result in imbalanced myosin expression, which could contribute to a disease-related phenotype, as mutations in *MYH6* have been shown to contribute to cardiomyopathies and congenital heart disease (Carniel et al. 2005; Ching et al. 2005).

## Discussion

In this study, we performed epigenetic and transcriptomic analysis on iPSC-CMs in various cell states. We identified a novel regulatory region that functions in cardiomyocytes and in response to critical extracellular stimuli. We developed a more comprehensive mechanism by which *MYH6* and *MYH7* are regulated, adding to previous work surrounding the C3 myosin enhancer (Gacita et al. 2021). We see evidence for varying activity, accessibility, and looping of CTCF-bound or proximal enhancers, R3, C3, and M3, across chambers of the heart and with various stimuli (GSK3i and ET-1) (**Fig. 3a, 6, Supplemental Fig. S2a, S5**). R3 had increased accessibility and a stronger signal for H3K27ac in atrial human samples than in ventricle samples (**Fig. 3a, Supplemental Fig. S2a**). Previous studies have shown that atrial chambers have higher *MYH6* expression than ventricle chambers (Locher et al. 2011; Gacita et al. 2021). As R3 activity declines, *MYH6* loops to a distal enhancer (M3), and MYH6 expression declines; *MYH6* eventually loops to more distant regulatory elements (**Fig. 4b, f, 6d-e, Supplemental Fig. S5**).

This CTCF-mediated looping is representative of a common occurrence within developmental gene regulation (Beagan et al. 2017; Kubo et al. 2021). We found that these enhancers play important roles in CM stress response and contain high-confidence CTCF motifs like other developmental enhancers (**Fig. 2e**) (Kubo et al. 2021; Cavalheiro et al. 2021; Beagan et al. 2017). Others have found that as maturation occurs, developmental CTCF enhancers become repressed, and genes begin to loop to other distal enhancers, which have downstream effects on gene transcription (Tsai et al. 2018; Cavalheiro et al. 2021; Beagan et al. 2017; Kubo et al. 2021). We find this to be the case with the R3 *MYH6* enhancer and find evidence for this occurring elsewhere in the genome in CMs (**Fig. 7**, **Fig. 2e**). This decline in enhancer activity is exacerbated by stress, resulting in an even larger decrease in proximal *MYH6* interactions and repression of *MYH6*.

We identified enrichment of genetic variation within R3 in patients with cardiomyopathies (**Fig. 8a**), including variants in a CTCF motif and a ZNF460/Zfx motif. We observed no such enrichment in regions R1 or R2. If the enriched variants impair R3 function, it would support the role of R3 in regulating *MYH6* expression and underscore the impact of genetic variation on complex human traits. Allelic imbalance of disease-related genes has been observed in patients with hypertrophic cardiomyopathy (HCM) and is proposed as a potential driving mechanism (Montag et al. 2017; Montag and Kraft). Studies investigating genetic variation within R3 could further contribute to this model. The activation of the R3 enhancer was antagonistic to the transcriptional alterations and functional differences occurring during culture with ET-1. Under stressed culture conditions R3-activated cells maintained higher *MYH6* expression, higher MYH6/C3 interactions, and did not observe a decline in calcium amplitude following culture with ET-1 (**Fig. 4f, 5g, 6g**). Alterations in calcium signaling were observed to occur before heart failure following ET-1 treatment in mouse models of heart disease (Mueller et al. 2011). *MYH6* dysregulation has been widely observed to influence heart disease progression, and *MYH6* reactivation has proven beneficial in countering heart disease phenotypes (Chen et al. 2021; Han et al. 2014; Ching et al. 2005; Locher et al. 2011; Herron et al. 2010). This work adds to the evidence supporting *MYH6* as a potential therapeutic target for treating heart failure. However, an *MYH6* gene therapy would be challenging due to the large size of the cDNA and the packaging limit of cardiac-tropic gene therapy vectors such as AAV. Thus, activating the endogenous *MYH6* through CRISPRa or other therapeutic modalities may be an appealing alternative (Yang et al. 2016). Therefore, studies of *in vivo MYH6* activation should be an area of future research. Our work establishes a foundational understanding of myosin regulation, providing a defined set of potential targets, experimental readouts, and anticipated outcomes for future *in vivo* studies.

Our study demonstrates the value of combining CRISPR-epigenome editing with chromatin conformation assays to functionally characterize regulatory elements and link them to their gene targets. We identified a key enhancer essential for maintaining precise state-specific myosin expression. We performed the first experiments looking at the effects of endogenous activation of *MYH6*, which revealed that altering myosin expression has pleiotropic effects in normal and stressed states. The epigenetic and transcriptomic profiling and functional data presented here highlight key enhancers important for cardiac stress response that may play key roles in heart disease progression. These enhancers may provide directions for developing future gene therapies for heart disease.

## Methods

### Plasmids

For CRISPRa experiments, we used the all-in-one lentiviral vector pLV hU6-gRNA hUbc-VP64-dSpCas9-VP64-T2a-Blast. The plasmid was generated from pLV hU6-gRNA(anti-sense) hUbC-VP64-dCas9-VP64-T2A-GFP (Addgene 66707), the GFP cassette was swapped out for a blasticidin resistance cassette. For CRISPRi experiments, the all-in-one lentiviral vector pLV hU6-gRNA hUbc-dSpCas9-KRAB-T2a-Puro was used (Addgene 71236). The gRNAs were cloned into their designated vector following the digestion of the plasmid with Esp3I (NEB) using T4 ligation methods (NEB) (**Supplemental Table S5**).

### Cell Lines

HEK293T cells were cultured in Dulbecco’s modified Eagle medium (DMEM) supplemented with GlutaMAX, 10% fetal bovine serum (FBS), 1 mM sodium pyruvate, 1× MEM non-essential amino acids, 10 mM HEPES, and 100 U/ml penicillin-streptomycin.

HL-1 cells were maintained on 0.02% gelatin (EMD Millipore) and 5 μg/ml fibronectin-coated plates, using Claycomb basal medium supplemented with 10% FBS, 0.1 mM norepinephrine (Sigma), 2 mM L-glutamine (EMD Millipore), and 100 U/ml penicillin-streptomycin(Claycomb et al. 1998). The human WTC11 iPSC cell line (UCSFi001-A) was maintained on Matrigel-treated (Corning) dishes in mTesR+ (Stemcell Tech).

### iPSC to cardiomyocyte differentiations

The cardiomyocyte differentiation protocol was adapted from a previously established small-molecule protocol (Lian et al. 2013). On day 0, cells were treated with 10 μM CHIR9902 (Cayman Chemical) in RPMI1640 (ThermoFisher) supplemented with B27 supplement without insulin (RPMI/B27-) (ThermoFisher). On day 2, the medium was replaced with RPMI/B27- and 5 μM IWP2 (VWR). On day 4, the media was changed to RPMI/B27-. From days 6 to 10, the medium was changed every 48 hours with RPMI1640 and B27 supplement (RPMI/B27+). From days 10 to 14, the medium was replaced every 48 hours with RPMI/B27+ without glucose, supplemented with ascorbic acid (Stemcell Tech), sodium DL-lactate (Sigma), and recombinant human albumin (Novus/R&D Systems) to purify cardiomyocytes. On day 14, iPSC-derived cardiomyocytes were dissociated using 0.05% Trypsin-EDTA (ThermoFisher) and replated on Matrigel-treated dishes. Following differentiation, cells were maintained in RPMI/B27+ with 100 U/ml penicillin-streptomycin. All differentiations were assessed to confirm successful cardiomyocyte differentiation.

### Immunofluorescent Staining and Imaging

iPSC-CMs were seeded into Matrigel-coated wells (Ibidi, 80826) at a density of 70k to 150k cells per cm². Twenty-four hours after seeding, cells were fixed in 4% PFA (VWR) in PBS for 15 minutes and washed three times in PBS containing 0.1% Triton-X 100 (Sigma, PBST). Cells were blocked and permeabilized in blocking solution containing PBST, 0.2M glycine, and 2.5% FBS for 1 hour. Primary antibodies against proteins of interest were applied for 1 hour. The following primary antibodies were used: Mouse anti-α-actinin (Sigma, 1:500) and Rabbit anti-phospho-histone H3 (pHH3, Cell Signaling Technology, 1:800). Cells were washed three times in PBST and then incubated with secondary antibodies and DAPI (ThermoFisher) for 50 minutes. The following secondary antibodies were used: Alexa Fluor 488 Goat anti-rabbit and Alexa Fluor 647 Donkey anti-mouse (ThermoFisher) (**Supplemental Table S5**). Finally, cells were washed three times in PBST and mounted with VECTASHIELD antifade mounting medium (VWR).

### Analysis of iPSC-CM Proliferation and Genomic Content

To assess the proliferative index and genomic content of iPSC-CMs, cells were differentiated as described above and allowed to mature for 1 week. Cells were cultured for an additional week with/without 4 μM CHIR9902 (GSK3i). During the final 48 hours, cells were exposed to 10 μM EdU (5-ethynyl-2’-deoxyuridine) (**Fig. 1a)**. Following treatment, cells were seeded for immunofluorescent imaging, fixed in 4% PFA the following day, and processed using the Click-iT Plus EdU kit with Alexa Fluor 555 (Invitrogen, ThermoFisher). Cells were then stained for pHH3 and α-actinin.

Cells were imaged using a Zeiss AxioObserver7 inverted microscope with a Zeiss Axiocam 305 color camera. Images were processed and analyzed using a combination of CellProfiler pipelines and R scripts (Stirling et al. 2021). The assistance of ChatGPT was used to generate the R script for the analysis of proliferation and cell cycle stage. CellProfiler was used to identify nuclei and measure the intensity of the signal for DAPI, EdU, and pHH3. R scripts were used for data wrangling, and GraphPad Prism was used for visualization and statistical analysis (**Supplemental Table S1, Fig. 1c-e**). EdU and pHH3 staining were used as markers of proliferation, while DAPI intensity was used to assess genomic content and determine cell cycle stage based on nuclear DNA content. Cell cycle distribution was estimated using genomic content across all cells analyzed (**Fig. 1**). This method has been used in combination with EdU incorporation to determine the cells actively proliferating or replicating DNA and those in a cell cycle arrest (Pereira et al. 2017).

### ATAC Sequencing

Nuclei preparation and sequencing analysis were based on methods previously developed in the lab (Corces et al. 2017; Cosgrove et al. 2024). A total of 1x10⁵ iPSC-CMs were harvested for each replicate and condition (**Supplemental Table S5**). ATAC sequencing libraries were sequenced on an Illumina NextSeq 2000 with paired-end 50 bp reads. Adapter sequences were trimmed, and reads were filtered by quality score using Trimmomatic (v0.32) (Bolger et al. 2014). Trimmed reads were aligned to hg38 using Bowtie (v1.2.3)(Langmead et al. 2009), allowing up to 2,000bp fragments (-X 2000), discarding multimapping reads (- m 1) and reporting best alignments (--best --strata). ENCODE blacklisted regions were removed using Picard MarkDuplicates and bedtools (v2.19.1) (Quinlan and Hall 2010). ATAC peaks were called using MACS2 (v2.1.1) callpeak function (Zhang et al. 2008). For visualization, deeptools bamCoverage was used to generate bigwig files with counts per million (CPMs) from deduplicated BAM files (Ramírez et al. 2014). Sample quality was assessed based on the number of uniquely mapped reads after blacklist removal. A union peak set was generated from MACS2 narrowPeak files of all samples. Count files for each sample were generated using featureCounts (**Supplemental Table S2**) (Liao et al. 2014). Differential accessibility within the union set of ATAC peaks was determined using DESeq2 (**Supplemental Table S2, Fig. 2a**) (Love et al. 2014). ATAC-seq profiles surrounding the MYH6 locus (**Fig. 2f-g**) were visualized in IGV (Robinson et al. 2011). The identification of TF binding motifs in differentially accessible regions was performed using MEME Suite (v5.5.7) (**Supplemental Table S2**)(Bailey et al. 2015b). All differential peaks with a *P_adj_* < 0.05 were considered for this analysis. Motifs with a q-value (FDR) < 0.1 and a TPM for the corresponding TF > 1 based on our RNA sequencing results were considered as potential TF binding sites and included in the analysis for **Fig. 2, 3**. Three rounds of analysis were conducted. First, motif discovery using the MEME Suite was performed on all differentially accessible peaks (*P_adj_* < 0.05) to identify general trends and TFs associated with peaks showing decreased accessibility (**Supplemental Table S2**). This analysis was repeated for peaks exhibiting increased accessibility (**Supplemental Table S2**). The results of this analysis are shown in **Fig. 2**. The final round of analysis focused specifically on region R3. These results, combined with genetic variants identified in participants from the UK Biobank, were used to assess the potential role of region R3 in cardiac health and disease. The findings are presented in **Fig. 3 and 7** (**Supplemental Table S2, Supplemental Fig. S5**). The assistance of ChatGPT was used to generate the R script for the analysis of the data generated using MEME Suite.

### RNA Sequencing

Total RNA was extracted from iPSC-CMs using the Norgen Total RNA Purification Plus Kit and sequenced by Azenta with standard RNA sequencing and rRNA depletion. Adapter sequences were trimmed, and reads were filtered by quality score using Trimmomatic (Bolger et al. 2014). Trimmed reads were aligned to hg38 using STAR (v2.4.1a) (Dobin et al. 2013). RPKM-normalized bedGraph files were generated using deepTools bamCoverage. UCSC bedGraphToBigWig utility was used to convert bedGraph files to bigwig format for genome browser visualization. Gene read counts were created using subread featureCounts (v1.4.6-p4) (**Supplemental Table S2**) (Liao et al. 2014). Differential expression analysis was performed using DESeq2, where counts are fit to a negative binomial general linearized model (GLM) and Wald statistics to determine the significance of differentially expressed genes (**Supplemental Table S2, Fig. 2b**) (Love et al. 2014). Visualization of RNA-seq profiles surrounding the MYH6 locus (**Fig. 2f-g**) was made in IGV (Robinson et al. 2011). Expression levels of TFs unique to differentially accessible regions of the genome were shown in (**Supplemental Fig. S3b-c**).

### Gene Ontology analysis using g:Profiler

Gene Ontology (GO) analysis was performed with g:Profiler and g:GOSt (v e11_eg58_p18_f463989d) on differentially expressed genes above a threshold for significant (*P_adj_*<0.01) and fold change in expression (|log_2_(FC)|>1) (Kolberg et al. 2023). A separate analysis was performed for genes decreasing or increasing in expression. All available annotation databases were used to assess pathway enrichment in differentially expressed genes; GO, KEG pathways, Reactome, CORUM, and others. Selected terms were presented in the main body of the text (**Fig. 2c-d**), and a complete list of results can be found in the supplemental data (**Supplemental Table S2**).

### Generation of UCSC browser tracks

All browser track visualizations were made using the UCSC Genome Browser for **Fig. 3**, and **all supplemental figures** (Nassar et al. 2023). The following are ENCODE accessions for the data used to generate **Supplemental Fig. S2a-c**, which can be found in **Supplemental Table S5** (Zhang et al. 2020; Abascal et al. 2020; Dunham et al. 2012).

For **Fig. 3** and **Supplemental Fig. S3 to S4,** the published data that was used or reanalyzed can be found in **Supplemental Table S5** (Zhang et al. 2020; Abascal et al. 2020; Dunham et al. 2012; Zhou et al. 2023; Akerberg et al. 2019; He et al. 2011). Visualization of the sample was for representation of H3K27ac in the human ventricle was noted by ENCODE to have lower read depth and mild bottlenecking. Mouse TEAD1 and NKX2.5 transcription factor ChIP-seq data were downloaded from GSE124008 and re-processed using the ENCODE ChIP-seq pipeline with default settings for transcription factor ChIP-seq; all TF ChIP-seq datasets from this dataset were processed, but only datasets with over 70% of reads from each replicate mapping to mm10 were visualized (Akerberg et al. 2019). Mouse p300 ChIP-seq data was downloaded from GSE195905 and reprocessed using the ENCODE transcription factor ChIP-seq pipeline with default settings for transcription factor ChIP-seq (Zhou et al. 2023). HL-1 cell transcription factor ChIP-seq data was downloaded from GSE21529 and reprocessed using the ENCODE transcription factor ChIP-seq pipeline with default settings for transcription factor ChIP-seq; apart from p300, only datasets with at least 20 million filtered reads at over 70% of reads mapping to mm10 were visualized (He et al. 2011). All visualized ChIP-seq datasets had NSC > 1.05 and RSC > 0.8. When visualizing transcription factor ChIP-seq data, fold-change bigwig files and IDR reproducible narrowpeak files were used; if multiple replicates were available, pooled reads from across all replicates were used for the bigwig files, and “conservative” IDR reproducible peak files were used for the narrowpeak files. Human cell-type-specific ATAC-seq signals from single-nucleus ATAC-seq data were taken from the Cardiac Atlas of Regulatory Elements (CARE) Portal (Hocker et al. 2021b). TF motifs displayed in the JASPAR2022 tracks are those for cardiac and other TFs, with motifs occurring more than expected by chance in the differential ATAC-seq motif analysis described above (Castro-Mondragon et al. 2022). Mouse regions R3 and M3 were defined as the 2kb region centered on the CTCF ChIP-seq signal from mouse heart samples. The mouse region C3 was previously defined (Gacita et al. 2021).

### Visualization of publicly available Hi-C data from the 3DIV database

Visualization of 3D chromatin conformation centered on the *MYH6* promoter and the enhancer region overlapping CMTM5 was generated using the 3DIV database (Jung et al. 2019; Yang et al. 2018). This database has analyzed about 400 publicly available Hi-C and promoter capture Hi-C datasets, including tissue samples from the left and right ventricles. We selected comparative interaction visualizations between the left and right ventricles using either the *MYH6* promoter or the myosin-specific enhancer region overlapping CMTM5 that we identified as the bait region. Regions in red indicate higher contact frequency in the left ventricle, while regions in blue indicate higher contact frequency in the right ventricle (**Supplemental Fig. S5**).

### Lentiviral Production, Delivery, and Antibiotic Selection

Lentiviral production was performed in HEK293T cells using Lipofectamine 3000 (Invitrogen), following previously described methods (McCutcheon et al. 2023b). The lentiviral supernatant was harvested 24 and 48 hours after transfection and centrifuged at 600g for 10 minutes to clear debris. The virus was then concentrated 100-fold using LentiX Concentrator (Takara Bio). iPSC-CMs and HL-1 cells were transduced at a multiplicity of infection (MOI) of approximately 0.3. Transduced cells were selected 48 hours post-transduction using puromycin (ThermoFisher) at 1 μg/mL for 4 days and blasticidin (ThermoFisher) at 2.5 μg/ml.

### Enhancer Activity via RT-qPCR Analysis

Cells were differentiated as described above and were left in culture for 2 weeks before use. iPSC-CMs were then transduced with lentiviral vectors expressing either dCas9^KRAB^ or ^VP64^dCas9 ^VP64^ along with a gRNA targeting the myosin enhancer or a non-targeting gRNA. After transduction, cells were enriched via antibiotic selection and grown for 15 days post-transduction. When used, ET-1 (Sigma) was added at a final concentration of 1 μM for the last 72 hours of culture. mRNA was extracted using the Norgen Total RNA Purification Plus Kit. Reverse transcription was performed using equal quantities of RNA for each sample with SuperScript VILO cDNA Synthesis. RT-qPCR was performed using equal amounts of corresponding cDNA per sample with Perfecta SYBR Green Fastmix (Quanta BioSciences) on the CFX96 Real-Time PCR Detection System (Bio-Rad). All primer products were verified via melt curve analysis. Relative expression levels were normalized to TBP and displayed as log2 fold change relative to controls. The significance for differences in gene expression was calculated on dCt values (normalized to *TBP*) using the designated statistical test with multiple hypothesis testing corrections. For HL-1 cells, similar protocols were followed. RNA was harvested 15 days post-transduction, with norepinephrine and FBS removed from specified samples for the last 48 hours of culture.

RT-qPCR primers for human *NPPA* (Hs.PT.58.4259173), *NPPB* (Hs.PT.58.19450190), mouse *Tbp* (Mm.PT.39a.22214839), *Myh6* (Mm.PT.58.31314128), *Myh7* (Mm.PT.58.17465550) were purchased from IDT. RT-qPCR primers for human *ATP2A2* and mouse *Atp2a2* came from previously published work (Traister et al. 2014; Angrisano et al. 2014). RT-qPCR primers for mouse *Nppa* and *Nppb* came from previously published work (Sergeeva et al. 2016b). RT-qPCR primers for mouse *Tnnt2* came from previously published work (Han et al. 2014). All RT-qPCR primer sequences can be found in (**Supplemental Table S5**).

### Analysis of cell size and genomic content

Cells were differentiated as described above to assess the cell size and genomic content of iPSC-CMs. iPSC-CMs were then transduced with lentiviral vectors expressing either ^VP64^dCas9 ^VP64^ along with a gRNA targeting the myosin enhancer or a non-targeting gRNA 25 days after the start of differentiation. After transduction, cells were enriched via antibiotic selection and replated on Matrigel-coated wells (Ibidi, 80826), as described above. For the last 48 hours of culture, cells were grown with RPMI/B27+ containing 1 μM ET-1 (stress) or vehicle (control). Cells were fixed in 4% PFA (VWR) in PBS for 15 minutes and washed three times in PBS containing 0.1% Triton-X 100 (Sigma, PBST) 20 days following transduction. Cells were blocked and permeabilized in blocking solution containing PBST, 0.2M glycine, and 2.5% FBS for 1 hour. Primary antibodies against proteins of interest were applied for 1 hour. The following primary antibody was used: Rabbit anti-TNNT2 (abcam, 1:500). Cells were washed three times in PBST and then incubated with secondary antibodies and DAPI (ThermoFisher) and Phalloidin-647 (ThermoFisher) targeting F-Actin for 50 minutes. The following secondary antibodies were used: Alexa Fluor 488 Goat anti-rabbit (ThermoFisher) (**Supplemental Table S5**). Finally, cells were washed three times in PBST and mounted with VECTASHIELD antifade mounting medium (VWR).

Cells were imaged using a Zeiss AxioObserver7 inverted microscope with a Zeiss Axiocam 305 color camera. Images were processed and analyzed using Fiji ImageJ (National Institutes of Health). The assistance of ChatGPT was used to generate the R script for **Supplemental Fig. S6b**. GraphPad Prism was used for visualization and statistical analysis (**Fig. 5b-c**). Statistical analysis was only performed across gRNA targets in the same media condition or across media conditions for an individual single gRNA condition, and a two-way ANOVA with Fisher’s LSD was used. The area covered by each individual cell was measured for only TNNT2-positive cells with defined borders. The total DAPI intensity measured within each cell area was used to measure genomic content.

### Flow cytometry to measure cell size of HL-1 cells

HL-1 cells were transduced with lentiviral vectors expressing ^VP64^dCas9^VP64^ and a gRNA targeting the myosin enhancer or a non-targeting gRNA. Cells were cultured for 21 days to allow for enhancer activation and gene expression alterations. Cells were dissociated and resuspended in flow buffer containing 1x PBS, 2 mM EDTA, and 0.5% bovine serum albumin. Cells were analyzed for size (FSC) and shape (SSC), gating for live and singlet populations (**Supplemental Fig. S6**). Median FSC values were used to determine differences in cell size between populations (**Fig. 5e**). A SH800 FACS Cell Sorter (Sony Biotechnology) was used for analysis.

### Calcium analysis for human iPSC-CMs

The iPSC-CMs used for the following experiments followed the same culture protocol as cells used to analyze cell size and genomic content following enhancer activation and stress. Imaging and analysis were performed across two separate differentiations and three separate transductions across each differentiation for each gRNA. For stress experiments, 24 hours prior to Ca2+ imaging, iPSC-CMs were incubated with either 1 μM ET-1 (stress) or vehicle (control) in RPMI/B27+. 24 hours following incubation, iPSC-CMs were washed once with 1X Ca2+-free Tyrode solution and then stained with 10 µM CAL-520 (ab171868, Abcam) for 1 hour. Following incubation, iPSC-CMs were then incubated with a 1:1 solution of RPMI/B27+ and 1X Tyrode with 1.8 mM CaCl2 for 15 minutes prior to imaging. iPSC-CMs were paced at 0.5Hz with an IonOptix MyoPacer field stimulator (IonOptix) for ten seconds. Zeiss Laser Scanning Confocal 510 Meta Microscope (Carl Zeiss AG) was used to capture line scans of iPSC-CMs at 0.1 μM per pixel.

Calcium imaging experiments were analyzed on Fiji ImageJ (National Institutes of Health) using the SparkMaster plug-in. SparkMaster was used to analyze Ca2+ transients and quantify sparks (Picht et al. 2007). SparkMaster features include spark duration, time to peak, width, amplitude, and tau. To quantify Ca2+ leak, multiply spark frequency with the average amplitude, width, and duration at half the maximum of all the sparks for the line scan of a cell to generate a total leak value. Regions of interest used to measure sparks and leaks were generated 2 seconds from the last paced beat. Statistical analysis was only performed across gRNA targets in the same media condition or across media conditions for an individual single gRNA condition. A two-way ANOVA with Fisher’s LSD was used.

### Analysis of Chromatin Looping at the *MYH6* locus by targeted enriched HiCAR

iPSC-CM and HL-1 were transduced with lentiviral vectors expressing ^VP64^dCas9^VP64^ and either a gRNA targeting the myosin enhancer or a non-targeting gRNA. Cells were enriched for transduction events using antibiotics and cultured for 12 days post-transduction. iPSC-CMs were treated with vehicle (water) or 1 μM ET-1 for the final 72 hours, while HL-1 cells were cultured in Claycomb medium without norepinephrine and FBS during the final 72 hours to promote normal gene expression. Live cells were dissociated 15 days post-transduction, centrifuged at 850g for 5 minutes, resuspended in 1 mL PBS, and fixed by adding 27.8 µL of 37% PFA for 10 minutes. The reaction was quenched with 80 µL of 2.5 M glycine, and the fixed cells were centrifuged and washed. HiCAR libraries were then prepared as previously described (**Supplemental Table S5**) (Wei et al. 2022).

HiCAR libraries were enriched for interactions related to the *MYH6/MYH7* locus using a biotin-probe pull-down. Target regions included the *MYH6* promoter and gene body, the *MYH7* promoter, M3, and other cardiac-specific gene promoters as controls (**Supplemental Table S3**). Biotinylated probes were ordered from Twist Bioscience, and their enrichment protocol was followed for HiCAR library enrichment. Universal blockers (Revvity, NOVA-5143231) were used during enrichment to prevent undesired pull-downs and noise from occurring in downstream analysis. The targeted region for human and mouse libraries covered approximately 60 kb of genomic space. Each library aimed for a sequencing depth of at least 15 million reads. HiCAR libraries were sequenced on an Illumina NextSeq 2000 with paired-end 50 bp reads.

Following sequencing, standard HiCAR analysis was performed using the pipeline from the original publication (Wei et al. 2022). Briefly, Raw fastq files were mapped to the reference genome by BWA-mem with the ‘–SP5M’ option. The aligned reads were processed with Pairtools (v1.1.0) to filter out the following low-quality reads: reads with MAPQ less than 10, non-rescuable chimeric reads, and self-ligated reads (Abdennur et al. 2024). Duplicated reads were also removed with Pairtools (Abdennur et al. 2024). 10kb bins were used to aggregate counts during the generation of the count tables. Visualization of these regions was generated by custom Python scripts. The visualization of reads across all samples was generated with HiGlass (v1.14.8) (**Supplemental Fig. S7**) (Kerpedjiev et al. 2018).

### Identifying significant changes in chromatin interactions

Genomic interactions were analyzed using the contact matrix generated by the HiCAR pipeline in DESeq2 (v1.42.0) with default parameters (**Supplemental Table S3**) (Love et al. 2014). Briefly, a 10kb resolution raw contact matrix was constructed within the following genomic range: chr14:23,303,865-23,451,365 in hg38 coordinates and chr14:55,111,190-55,258,690 in mm39 coordinates. Those contact matrices were used as input for the differential test. All replicates were considered for each condition during the test. The Wald test was performed to assess the statistical significance of genomic pairs between the tested conditions. Benjamin-Hochberg test was performed to correct multiple hypothesis testing with 0.05 as a false discovery rate cutoff for the significance of the adjusted p-value.

### Analysis of genetic variants from UKBiobank samples within the myosin-specific enhancer

UK Biobank DRAGEN population level whole genome sequencing (WGS) data (500k release) was analyzed to identify variants within regions R1 to E3 associated with the myosin-specific enhancer (chr14:23377559-23378416 for region R1, chr14:23378863-23379551 for region R2, and chr14:23379863-23380961 for region R3, hg38) (Bycroft et al. 2018). Full cohort description and WGS processes are described elsewhere (Hofmeister et al. 2023). Briefly, WGS data (30x) was re-processed using the DRAGEN 3.7.8 pipeline for both individual and joint-called data for all participants (Scheffler et al.). In total, there were 12,928 participants with cardiomyopathies and 473,770 controls. Individuals were included in the cardiomyopathies group based on the criteria provided in the supplement and did not have coronary artery disease (CAD) (**Supplemental Table S4**). Individuals were considered a control when they did not meet the criteria for cardiomyopathies and did not have CAD (**Supplemental Table S4**). VCFtools (version 0.1.16) was used to filter for variants present within each region in participants with cardiomyopathy (n=4320 defined using self-report, electronic health record, and cardiac MRI data available in the UK Biobank database (Danecek et al. 2011). Likewise, variants within this region were filtered for in individuals without cardiomyopathy (controls; n=192,805). Specific criteria for defining CM participants and controls are available in (**Supplemental Table S4**). A total of 63 variants were identified in CM participants across intronic and noncoding regions overlapping the coordinates mentioned above (**Supplemental Table S4, Fig. 3b, 8a**). Fisher’s exact tests were performed to compare variant frequencies in cardiomyopathy participants to controls for each of the 31 variants identified in CM participants. Variant tolerance for variants within region R3 was assessed using JASPAR transcription factor binding profiles, evaluating the impact of each variant on transcription factor binding motifs. The tolerance score was calculated as ((Alt_value - Ref_value) / Total_value), with possible scores ranging from -1 to 1. A score of 0 indicates no effect, 1 represents a de novo binding motif, and -1 indicates a maximal decrease in base tolerance for the given variant.

Variants found across all regions were scored for their potential pathogenic or disruptive effects using Combined Annotation-Dependent Depletion (CADD v1.7) (Schubach et al. 2024). Those scores can be found in the supplemental data (**Fig. 8a, Supplemental Table S4**).

## Supporting information

GEO accession numbers

## Declarations

### Ethics approval and consent to participate

Not applicable

### Consent for publication

Not applicable

### Availability of data and materials

All code related to iPSC-CM proliferation, ATAC-seq, RNA-seq, motif analysis, and HiCAR analysis is available at https://github.com/Gersbachlab-Bioinformatics/myosin_enhancer. All code for the following assays can be found at the indicated link:

iPSC-CM proliferation analysis: https://github.com/Gersbachlab-Bioinformatics/myosin_enhancer/tree/main/figure1_scripts

ATAC-seq and RNA-seq analysis: https://github.com/Gersbachlab-Bioinformatics/myosin_enhancer/tree/main/figure2_scripts

HiCAR pipeline and downstream analysis: https://github.com/Gersbachlab-Bioinformatics/myosin_enhancer/tree/main/figure5_scripts

The ENCODE ChIP-seq pipeline https://github.com/ENCODE-DCC/chip-seq-pipeline2.

All raw sequencing data and analysis files for ATAC-seq, RNA-seq, and HiCAR can be found at https://www.ncbi.nlm.nih.gov/geo/. The accession ID for those files are: GSE283424, GSE283426, GSE283427, GSE283430, and GSE283432. The datasets supporting the conclusions of this article are included within the supplemental data.

All data generated or analyzed during this study are included in this article or are available from the corresponding author on reasonable request.

### Authors’ contributions

TA, RK, and CAG conceived the study. TA and CAG wrote the paper with the help of all authors. TA was primarily responsible for the design of all experiments, performed all iPSC to CM differentiations, generated virus for gene editing, processed all samples for data generation, and led analysis and interpretation of the generated data. IMK reprocessed publicly available data related to epigenetic profiling, generated the associated figures, assisted in their interpretation, and assisted in preparing plasmid and lentivirus. KTH assisted in the production of lentivirus used in these studies. TA, BC, IJ, YD, and CAG participated in the design, analysis, and interpretation of the HiCAR experiments. TA, RMP, ED, APL, and CAG participated in the design, analysis, and interpretation of the calcium imaging experiments. DT prepared all HiCAR libraries for NGS. MER and SS contributed to the design and analysis of the genetic variants in the UK Biobank. All authors contributed to data interpretation and writing of the manuscript.

## Acknowledgments

We thank the Duke Sequencing Core and Duke Cell Culture Facility for assistance. We also thank the teams at the High-throughput Applied Research Data Analysis Cluster (HARDAC) and Duke Computing Cluster (DCC) for computing resources. Schematics were created with BioRender.com.

