## Supplementary Figures S1-S7 for "A gene regulatory element modulates myosin expression and controls cardiomyocyte response to stress"

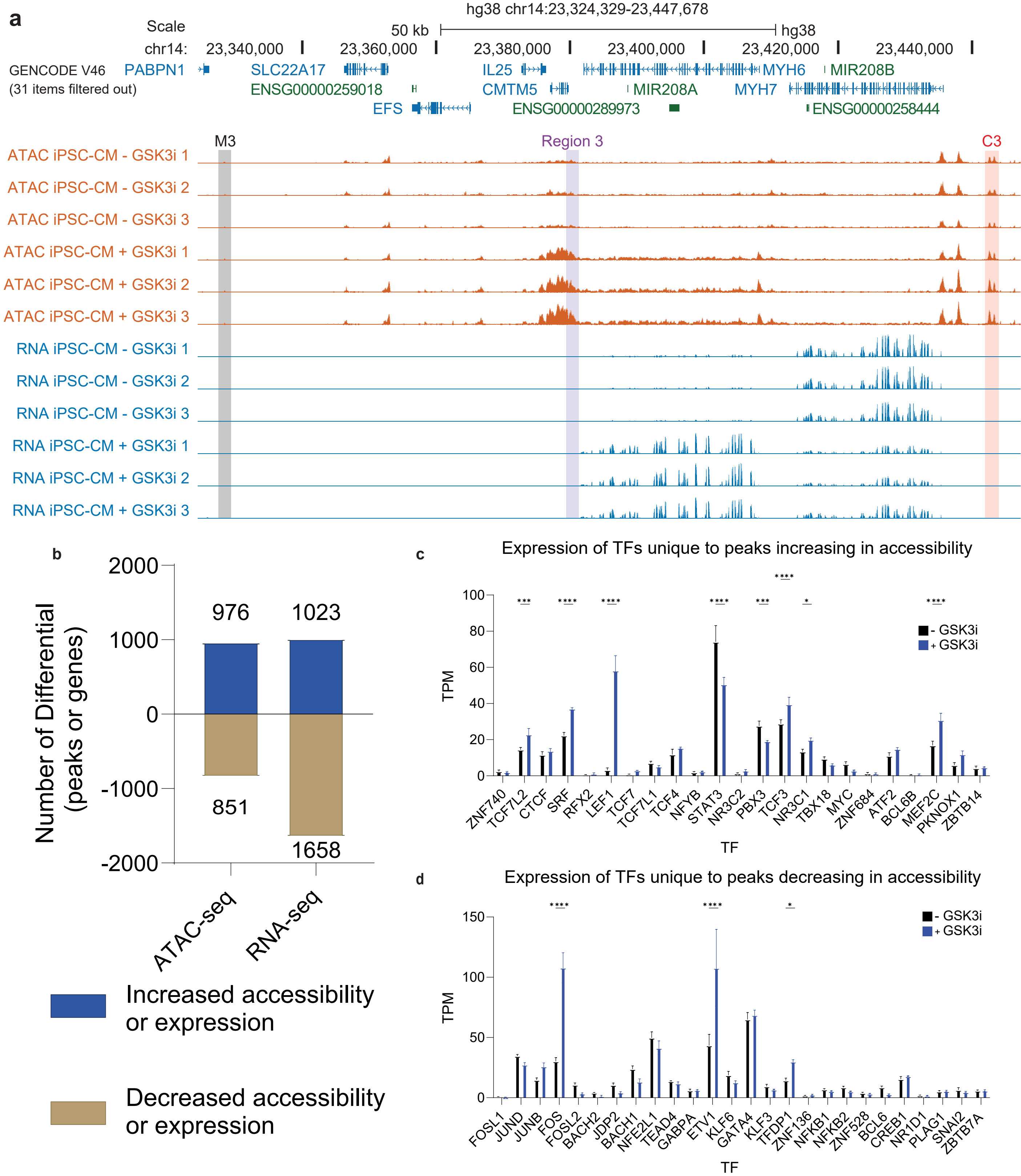


**Fig. S1 GSK3 inhibition reproducibly alters gene expression and chromatin accessibility**

a) Visualization of ATAC-seq (orange) and RNA-seq (blue) profiles surrounding the MYH6 locus. GSK3 inhibition leads to a reproducible increase in chromatin accessibility overlapping the cCRE region 3' of the MYH6 gene body. Additionally, there is a consistent shift in gene expression from MYH7 to MYH6 following GSK3i treatment. **b**) Number of differentially accessible ATAC peaks and differentially expressed genes following GSK3 inhibition. **c-d**) TPM of transcription factors with motifs unique to ATAC peaks, showing increased (**c**) or decreased (**d**) accessibility. Only motifs with an FDR < 0.1 were considered. See also Fig. 2 and Additional file 3.

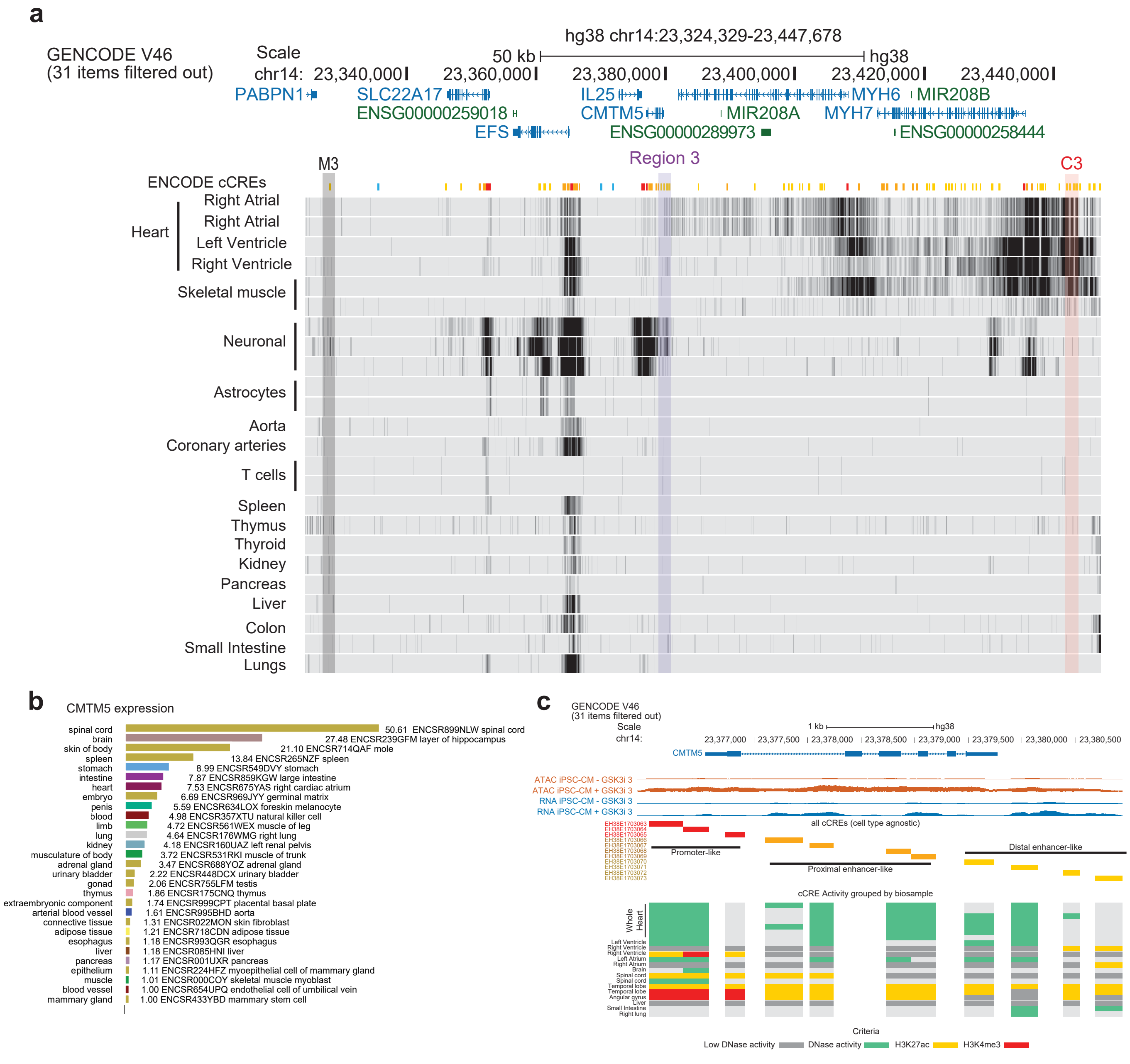


**Fig. S2 ENCODE cCRE activity across assorted human tissue types**

**a)** Visualization of ENCODE H3K27ac data around the MYH6 locus in various human tissues. **b)** Expression levels of CMTM5 across ENCODE tissue and primary cell samples, shown as TPM + 0.01. **c**) Visualization of ENCODE data indicating cCRE (enhancer) activity across selected tissue types. The first four rows show tracks generated from our ATAC-seq (orange) and RNA-seq (blue) data. Following these, ENCODE cCREs overlapping Regions R1-R3 are displayed with their classifications. These enhancers overlap RNA-seq signals in our iPSC-CM, which increase after GSK3i treatment. Lastly, the summary of cCRE activity supporting cCRE activity across various tissue types. See also Fig. 3

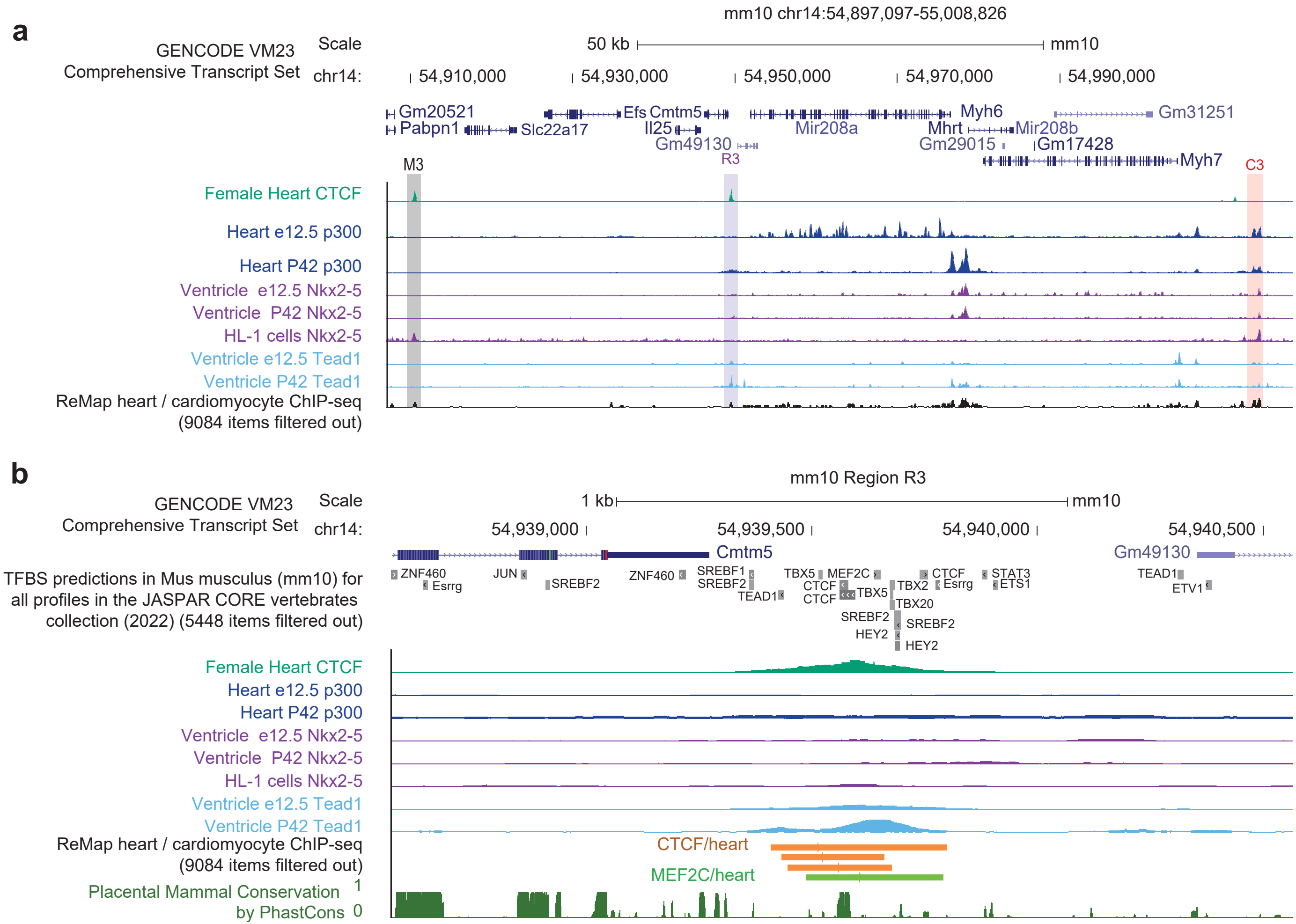


**Fig. S3 Epigenetic landscape surrounding a cardiac-specific regulatory element in mouse samples**

**a-b**) Visualization of epigenetic datasets surrounding the Myh6 locus (**a**) and Region R3 (**b**). Data include CTCF-ChIP (green), p300-ChIP (dark blue), Nkx2-5 (purple), Tead1 (light blue), and ReMap ChIP-seq data from cardiac-related studies. The cCRE regions are highlighted as follows: M3 (grey), R3 (purple), and C3 (orange). See also Fig. 3.

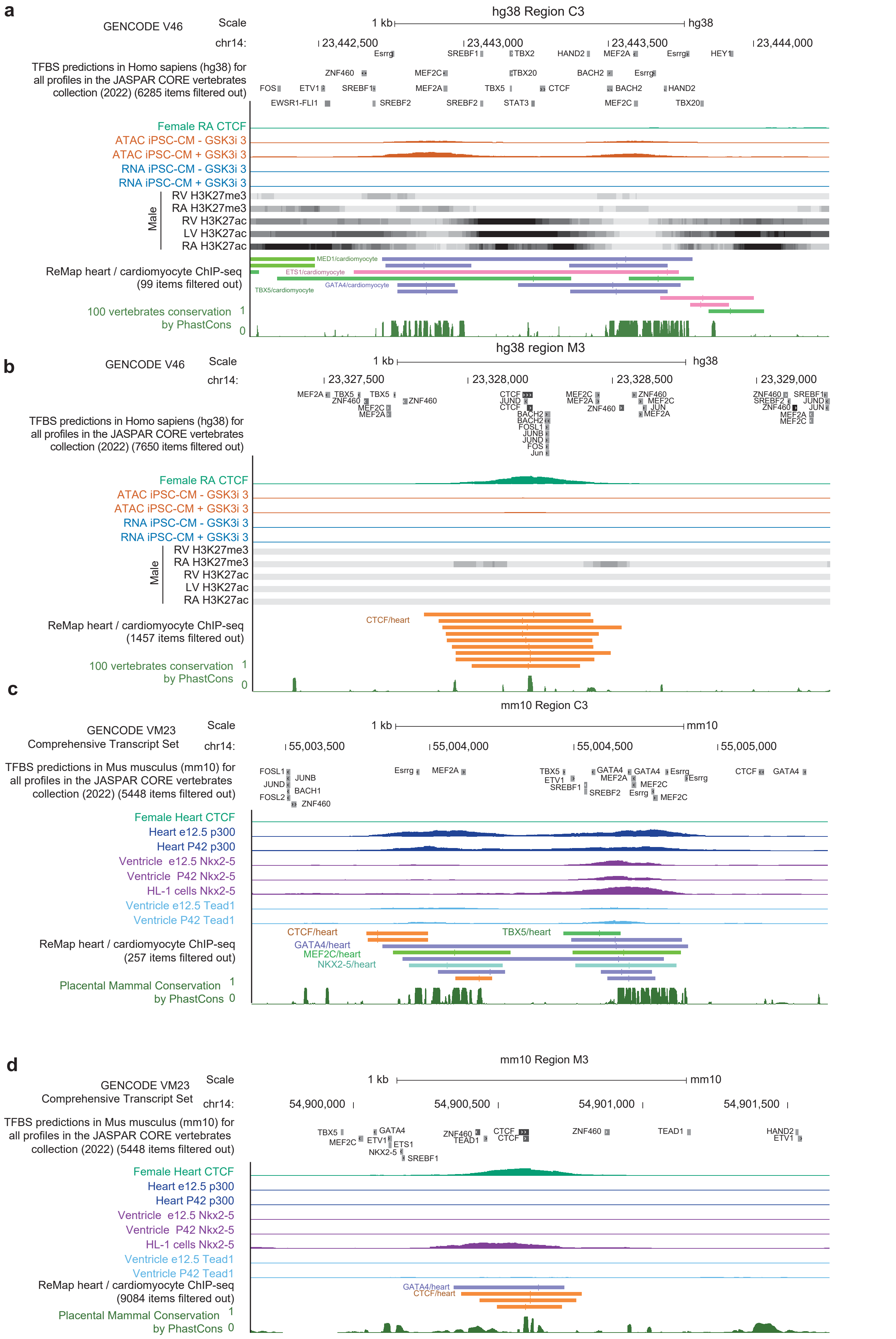


**Fig. S4 Epigenetic landscape surrounding the distal myosin enhancers C3 and M3 in humans and mice**

**a-b**) Visualization of epigenetic datasets surrounding Region C3 (**a**) and M3 (**b**). Datasets include CTCF-ChIP (green), ATAC-seq profiles in iPSC-CMs and single-cells ATAC-seq human hearts (orange), RNA-seq profiles in iPSC-CMs (blue), histone ChIP-seq density plots, and ReMap ChIP-seq data from cardiac-related studies.
**c-d**) Visualization of Epigenetic Datasets Surrounding Region C3 (**c**) and M3 (**d**). Data include CTCF-ChIP (green), p300-ChIP (dark blue), Nkx2-5 (purple), Tead1 (light blue), and ReMap ChIP-seq data from cardiac-related studies. The cCRE regions are highlighted as follows: M3 (grey), R3 (purple), and C3 (orange). See also Fig. 3.

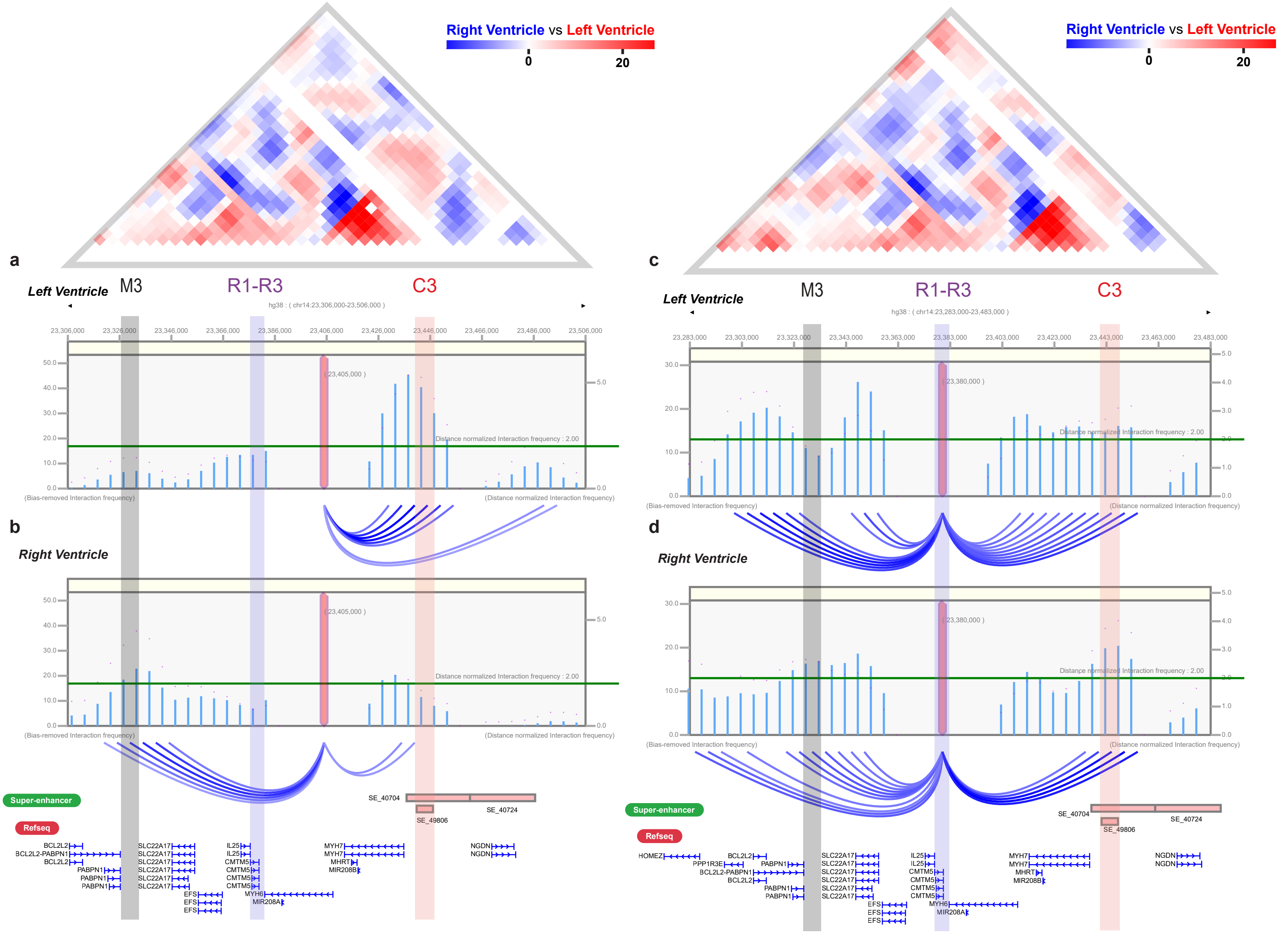


**Fig. S5 Chromatin looping data for the MYH6 promoter and R1-R3 in the Left and Right ventricle chambers of the heart.**
**a-b**) Visualization of chromatin looping between the promoter of MYH6 and the surrounding genomic environment in the left ventricle chamber **a**) and the right ventricle chamber **b**). In the left ventricle, MYH6 predominantly loops to the C3 enhancer, while in the right ventricle, MYH6 predominantly loops to the M3 enhancer. **c-d**) Visualization of chromatin looping between the promoter of MYH6 and the surrounding genomic environment in the left ventricle chamber **c**) and the right ventricle chamber **d**). In both chambers,R1-3 loops in both directions; however, R1-R3 exhibits fewer interactions with MYH6 in the right ventricle chamber compared to the left ventricle chamber. See also Fig. 3.

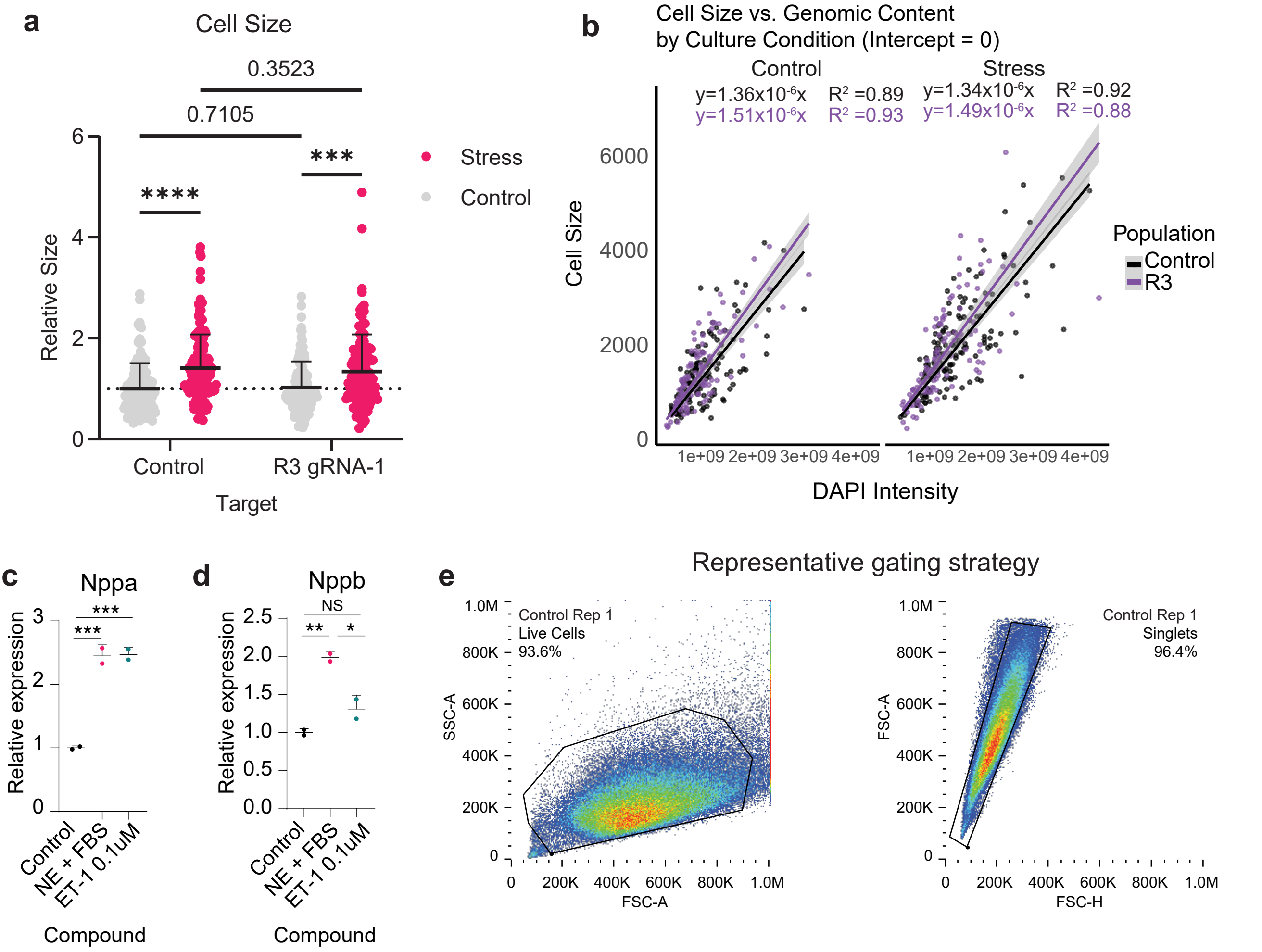


**Fig. S6 Analysis of functional measurements of cellular response to stress**
**a**) Normalized cell size for all cells measured of human iPSC-CMs represented 20 days following activation of each designated target ± ET-1 1μM for the last 48 hours of culture, n = 3 replicates. Statistics were calculated with a two-way ANOVA with Fisher's LSD. **b**) Visualization of individual cell size vs the genomic content (DAPI Intensity) within each cell separated by culture condition (Control = left, Stress = right) and target (Control = black, R3 = purple). Formulas for each linear regression with corresponding $R^2$ values are displayed. **c-d**) RT-qPCR analysis of Nppa (**c**) and Nppb (**d**) expression in HL-1 mouse atrial cardiomyocytes 48 hours after treatment with the indicated compounds. Relative expression is normalized to TBP and control (n = 2 replicates, mean ± SD). Statistical comparisons were made using dCt values (normalized to TBP) and one-way ANOVA with Tukey's post hoc test. Significant Padj values: Nppa: Control vs. NE+FBS ($P_{adj}$ = 0.0009), Control vs. ET-1 ($P_{adj}$ = 0.009), NE+FBS vs. ET-1 ($P_{adj}$ = 0.9828); Nppb: Control vs. NE+FBS ($P_{adj}$ = 0.0089), Control vs. ET-1 ($P_{adj}$ = 0.1104), NE+FBS vs. ET-1 ($P_{adj}$ = 0.0346). **e**) Gating strategy for cell size measurement: live cells were gated based on FSC-A and SSC-A, followed by a singlet gate to isolate the subpopulation of living cells, and median FSC-A per replicate was measured. See also Fig. 5.

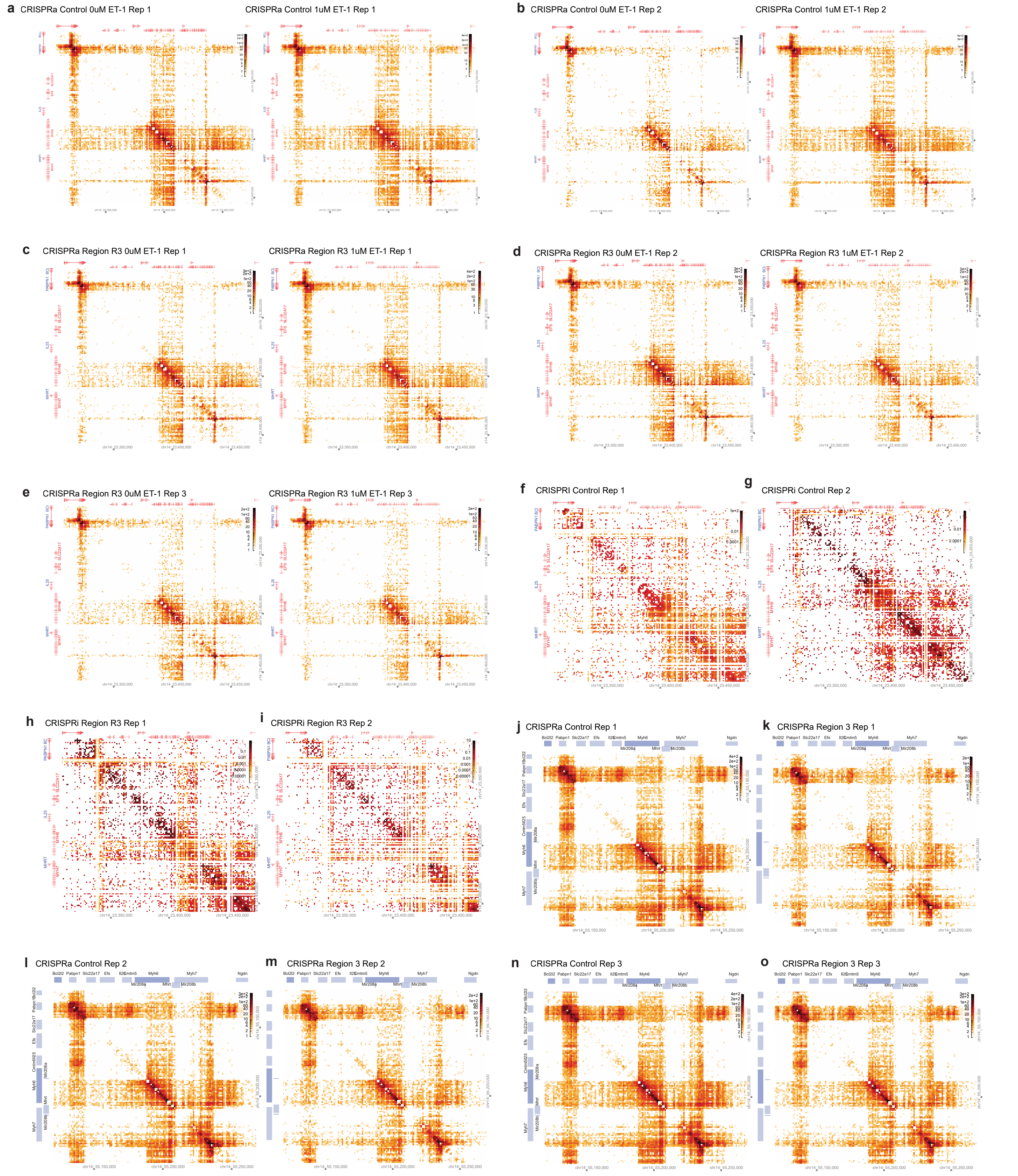


**Fig. S7 Visualization of targeted enrichment of HiCAR libraries in iPSC-CMs and HL-1 cells**

**a-b**) Contact heatmaps (hg38) at 1kb bin resolution, illustrating targeted enrichment of HiCAR libraries in iPSC-CMs transduced with $^{VP64}$dCas9$^{VP64}$ + Control gRNA +/- ET1 treatment for 72hrs. **a**) Replicate 1. **b**) Replicate 2. **c-e**) Contact heatmaps (hg38) at 1kb bin resolution, illustrating targeted enrichment of HiCAR libraries in iPSC-CMs transduced with $^{VP64}$dCas9$^{VP64}$ + Region R3 gRNA-1 ± ET1 treatment for 72hrs. **c**) Replicate 1. **d**) Replicate 2. **e**) Replicate 3. **f-i**) Contact heatmaps (hg38) at 1 kb bin resolution, illustrating targeted enrichment of HiCAR libraries in iPSC-CMs transduced with dCas9KRAB + gRNA. **f**) Control gRNA replicate 1. **g**) Control gRNA replicate 2. **h**) Control gRNA replicate 1. **i**) R3 gRNA replicate 2. **j-o**) Contact heatmaps (mm39) at 1 kb bin resolution, illustrating targeted enrichment of HiCAR libraries in HL-1 cells transduced with VP64dCas9VP64 + gRNA. **j,l,n**) Control gRNA replicates 1 to 3. **k,m,o**) Region R3 gRNA-2 replicates 1 to 3. See also Fig. 6, Additional file 6.
